# Defining the Cell Surface Cysteinome using Two-step Enrichment Proteomics

**DOI:** 10.1101/2023.10.17.562832

**Authors:** Tianyang Yan, Lisa M. Boatner, Liujuan Cui, Peter Tontonoz, Keriann M. Backus

**Author notes:** Corresponding Author: Keriann M. Backus, Biological Chemistry Department, David Geffen School of Medicine, UCLA, Los Angeles, CA, 90095, USA,.

## Abstract

The plasma membrane proteome is a rich resource of functional and therapeutically relevant protein targets. Distinguished by high hydrophobicity, heavy glycosylation, disulfide-rich sequences, and low overall abundance, the cell surface proteome remains undersampled in established proteomic pipelines, including our own cysteine chemoproteomics platforms. Here we paired cell surface glycoprotein capture with cysteine chemoproteomics to establish a two-stage enrichment method that enables chemoproteomic profiling of cell <u>Surf</u>ace <u>Cys</u>teinome. Our “Cys-Surf” platform captures >2,800 total membrane protein cysteines in 1,046 proteins, including 1,907 residues not previously captured by bulk proteomic analysis. By pairing Cys-Surf with an isotopic chemoproteomic readout, we uncovered 821 total ligandable cysteines, including known and novel sites. Cys-Surf also robustly delineates redox-sensitive cysteines, including cysteines prone to activation-dependent changes to cysteine oxidation state and residues sensitive to addition of exogenous reductants. Exemplifying the capacity of Cys-Surf to delineate functionally important cysteines, we identified a redox sensitive cysteine in the low-density lipoprotein receptor (LDLR) that impacts both the protein localization and uptake of LDL particles. Taken together, the Cys-Surf platform, distinguished by its two-stage enrichment paradigm, represents a tailored approach to delineate the functional and therapeutic potential of the plasma membrane cysteinome.

## Introduction

The cell surface proteome regulates nearly all aspects of cellular function, spanning, cell-cell communication^1–3^, signal transduction^4–6^, and changes to cell state^7^, including activation^8–10^, proliferation^11,12^, and death^13^. Consequently nearly 50% of modern drug targets are found in the membrane proteome, which is also rich in biomarkers indicative of disease^14–18^. Notably, over 200 cell surface proteins have been reported as overexpressed in human cancers^19^. Despite their prominence as high value therapeutic targets, the hydrophobicity and low abundance of most cell surface proteins complicates functional characterization and therapeutic targeting of this important fraction of the proteome^20–23^.

A number of high value biochemical and mass spectrometry based proteomic techniques have been developed to improve capture and identification of the cell surface proteome, most notably cell surface biotinylation^24–29^ and cell surface capture (CSC)^30–37^. As the N-hydroxysuccinimidobiotin (NHS-biotin) reagent used in cell surface biotinylation has been shown to label intracellular proteins^38,39^, CSC has emerged as a favored method for specifically enriching membrane bound glycoproteins. CSC leverages the heavy glycosylation of nearly all membrane proteins—90% of the proteins on the cell surface have been reported to be glycosylated^40^—to afford high specificity capture of the cell surface proteome. In the CSC method, cis-diols found on all glycans are first oxidized to aldehyde groups in the presence of sodium periodate. Subsequently, these aldehyde moieties are trapped with aminooxy- or hydrazide-biotin reagents, which affords robust biotinylation of cell surface proteins, via formation of a stable oxime or hydrozone. Biotinylated protein groups are then identified via established enrichment-based proteomic workflows. Showcasing the widespread utility and adoption of the CSC, recent studies have implemented this chemistry to identify and quantify surfaceome alterations in cancer cell lines^18,41–44^, such as overexpression of CD166 in head and neck squamous cell carcinoma^45^ and carbonic anhydrase 12 in colon cancer^46^.

Cysteine residues are a ubiquitous, functionally important, and therapeutically relevant feature of cell surface proteins^47^. Membrane proteins are particularly rich in structural disulfides and redox-active cysteines are widely known to regulate membrane protein function^48–52^. Multiple cell surface cysteines are reported to be redox active, including C53 of aquaporin-8 AQP8, persulfidation of which is known to gate H_2_O_2_ to regulate cell stress^53^ and C23, C45, and C106 of high mobility group box 1 protein HMGB1, reduction of which are known to be crucial for anti-inflammatory processes^54,55^. Activation-induced changes to cysteine oxidation have been linked to enhanced T cell activitation^56–58^ and cysteines on CD4 and GP120 have been implicated in the entry of HIV in T cells and in the regulation of HIV infection of T cells^59,60^. Cell surface cysteines have also been targeted by drugs and drug-like molecules, as showcased by Afatinib and Ibrutinib, blockbuster anti-cancer agents, that function by selectively and irreversibly labeling conserved non-catalytic cysteines found within kinase active sites^61,62^. Consequently, identifying redox sensitive and potentially druggable cysteine residues in membrane proteins is of high value for expanding the scope of the druggable proteome.

Mass spectrometry-based cysteine chemoproteomics is one such method that is well positioned to pinpoint such residues in membrane proteins. As shown by previous reports^63–67^, including our own, the human proteome harbors thousands of cysteines amenable to modification by druglike molecules—each cysteine-ligand pair represents the potential starting point for a drug development campaign. Similar to small molecule screening platforms, redox-directed chemoproteomic methods, including biotin-switch^68^, OxICAT^69^, SP3-Rox^70^, QTRP^71^, and Oximouse^72^, robustly report the relative and absolute oxidation states of cysteines analyzed from bulk proteomes. By pairing these methods with proximity labeling, recent methods including our own Cys-LOx have begun to shed light on subcellular compartment-specific cysteine oxidation states, which can vary markedly depending on protein localization^73,74^. Whether analogous approaches can be extended to the cell surface proteome remains to be seen.

Here, we establish cell <u>Surf</u>ace <u>Cys</u>teine enrichment (Cys-Surf), which pairs cell surface capture (CSC) with cysteine chemoproteomics to reveal the redox and ligandable cell surface cysteinome. Cys-Surf analysis of Jurkat and primary T cells identified 2,836 reactive cell surface cysteines on 1,046 proteins, which represents a >4-fold improvement of cell surface specificity as compared to bulk proteome analysis. Of these residues, 821 were not previously identified by cysteine chemoproteomics as reported by CysDB^63^, and we find that 211 proteins are ligandable by cysteine reactive compounds. By pairing Cys-Surf with our previously reported cysteine redox proteomic method SP3-Rox^70^, we quantify the absolute oxidation states of 1,246 cell surface cysteines on 489 proteins, reveal marked cysteine reduction during T cell activation, and stratify membrane cysteines sensitive to addition of exogenous oxidants and reductants. Guided by this Cys-Surf analysis we pinpoint a heretofore uncharacterized reduction-sensitive cysteine in an EGF repeat of the LDLR receptor, which impacts receptor localization and lipoprotein uptake.

## Results

### Establishing an SP3-enabled aminooxy-biotin cell surface platform

To establish our cell Surface Cysteine enrichment platform (Cys-Surf) (**Figure 1A**), we envisioned pairing cell surface capture (CSC)^30,31^ with cysteine peptide enrichment. Guided by our prior study, which paired TurboID proximity labeling with similar cysteine peptide enrichment to achieve subcellular redox proteomics^73^, we expected that efficient protein recovery throughout the Cys-Surf workflow would be essential to achieve high cysteine coverage.

**Figure 1.**
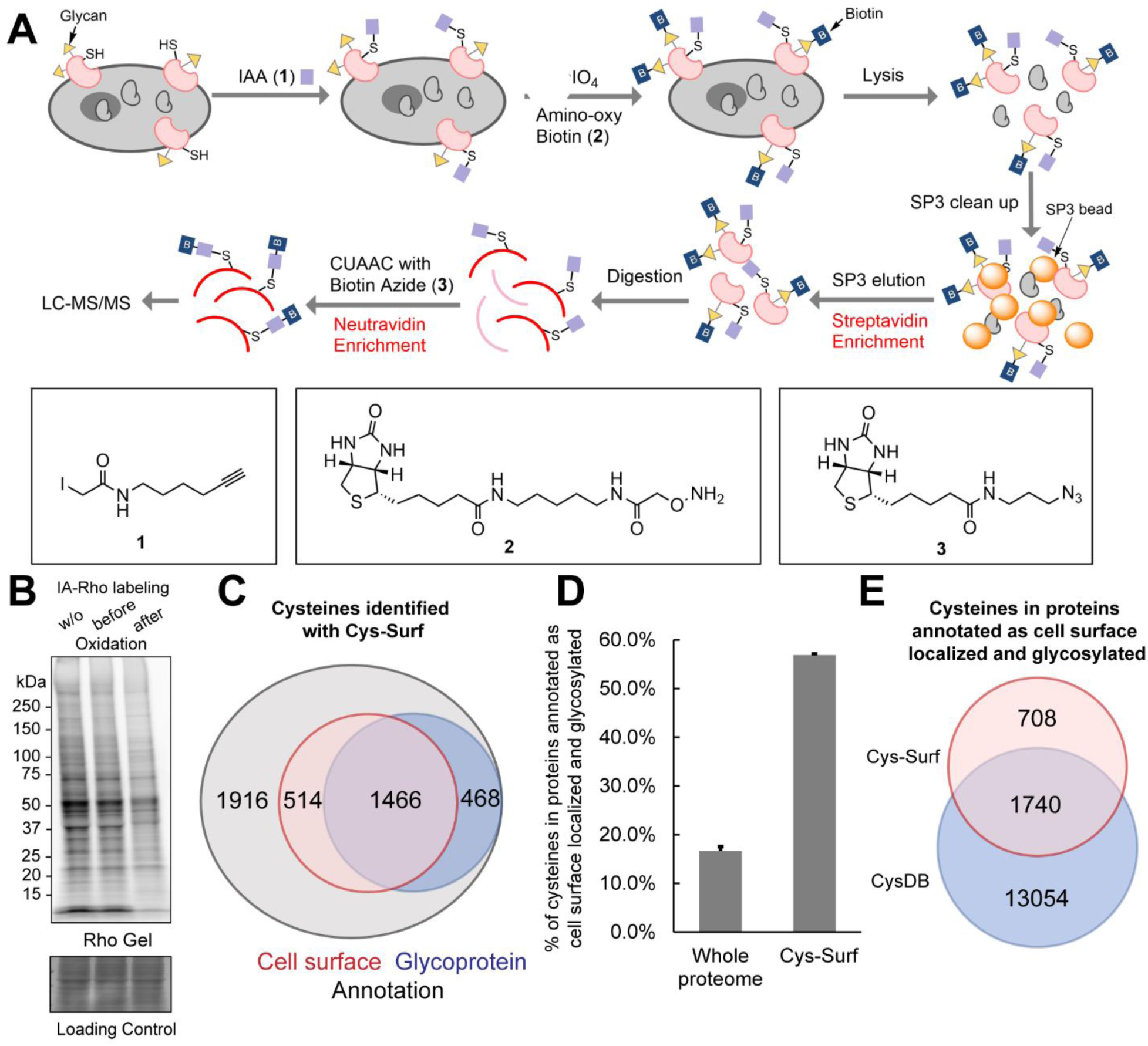
Cell surface cysteine enrichment enabled with cell surface capture and two-step biotinylation. **A)** The schematic workflow of Cys-Surf. **B)** Iodoacetamide-Rhodamine (IA-Rho) gel of Jurkats cells labeled with 5 μM IA-Rho without oxidation or before and after oxidation with NaIO_4_. **C)** Venn diagram of cysteines identified with Cys-Surf annotated as on the cell surface or from glycoproteins. **D)** Percentage of cysteines in proteins annotated as cell surface localized and glycosylated identified from MS analyses of trypsin digests from whole-cell and with Cys-Surf. Data is represented as mean ± stdev. **E)** Venn diagram of cysteines in proteins annotated as cell surface localized and glycosylated identified with Cys-Surf and in CysDB. MS experiments were conducted in 3 replicates in Jurkats cells. All data can be found in **Table S1**.

Therefore, we first prioritized establishing and enhancing the CSC portion of our method. We observed saturable labeling for aminooxy-biotin, with maximal labeling achieved with 1 mM treatment conditions (**Figure S1A**). As we had previously found that the single-pot, solid-phase-enhanced sample preparation (SP3) method^75^ afforded enhanced recovery of biotinylated samples^64,76^, we opted to incorporate SP3 into the CSC workflow. We established an SP3-based protein cleanup method in which the SP3 resin is employed to decontaminate samples prior to protein enrichment on streptavidin resin (**Figure S1B**). By comparing wash and protein elution conditions, we found that EtOH outperformed MeCN as the washing solvent and that addition of detergent (e.g. 0.2% SDS, which is not required for peptide elution^64,77^) is essential for protein recovery from SP3 resin (**Figure S1C and S1D**). Addition of SP3 to the CSC method afforded a modest yet significant improvement in peptide coverage (**Figure S1E**). As further confirmation that our implementation of CSC was robustly capturing cell surface glycoproteins, we additionally subjected enriched peptides to PNGase F digest (**Figure S2A**). We used the unique mass shift (0.98 Da) that remains on glycosylated asparagine residues after digest, to identify the glycosylation sites. ∼53% of detected peptides feature putative glycosylation sites and, of these, ∼93% belong to proteins annotated as localized to the cell surface (**Figure S2B**).

### Establishing cell <u>Surf</u>ace <u>Cys</u>teine enrichment (Cys-Surf)

We next paired our established CSC platform with cysteine labeling to establish our Cys-Surf workflow shown in **Figure 1A**. We expected that the sodium periodate used to oxidize glycans in the CSC workflow likely would also oxidize exposed cysteine thiols. Confirming this likelihood, we observed a decrease in cysteine reactivity towards iodoacetamide rhodamine (IA-Rho) following treatment of with sodium periodate; in contrast the IA-Rho signal remained unchanged when sodium periodate was applied after cysteine capping (**Figure 1B**). To circumvent this oxidation, we opted to first cap all cysteines in situ with the pan-cysteine reactive probe iodoacetamide alkyne (IAA;**1**).

The subsequent two-stage Cys-Surf capture workflow then proceeded smoothly. Following streptavidin-based CSC and on-resin digestion, cell surface peptides were collected and subjected to peptide-level biotinylation via click conjugation. After a second round of enrichment on neutravidin resin, the biotinylated peptides were then analyzed by liquid chromatography tandem mass spectrometry (LC-MS/MS). In aggregate, Cys-Surf analysis of Jurkat Cells captured a total of 4,364 unique cysteines in 1,889 total proteins. This coverage compares favorably to our previously reported two-stage enrichment platform using TurboID, in which we typically captured 500-1,500 total cysteines found on 300-800 proteins^73^.

To facilitate evaluation of the cell surface specificity of the Cys-Surf platform, specifically the fraction of captured cysteines that are localized to cell surface proteins, we generated a comprehensive cell surface protein database aggregating the localization information from the Human Protein Atlas^78^, UniProtKB^79^ and CellWhere^80^ (**Table S1**). In aggregate, 99,811 cysteines found in 7,018 proteins were annotated as localized on the cell surface by one or more databases analyzed. Consistent with our previous study in which we performed similar localization analysis for intracellular compartments^81^, we observed that a substantial fraction of the proteins with cell surface localization are also annotated as localizing to one or more additional subcellular compartments (4,339 total). As these multi-localized proteins could confound our analyses, we generated an additional dataset of glycosylated proteins generated from UniProtKB annotations as orthogonal evaluation of the method, which includes 85,933 cysteines on 5,139 proteins (**Table S1**). In total, 3,929 proteins are shared between both datasets.

Overall, Cys-Surf identified 1,980 cysteines in 700 proteins with annotated cell surface localization and 1,934 cysteines in 633 proteins annotated as glycoproteins (**Figure 1C** **and S3**). ∼75% of the glycoprotein cysteines (1,466/1,934) are also annotated as cell surface cysteines (1,466/1,980) (**Figure 1C**), which is consistent with the CSC enrichment of glycoproteins on the cell surface. Altogether, 56% (2,448/4,364) of cysteines identified with Cys-Surf are annotated either on the cell surface or in glycoproteins. By comparison, only ∼15% of cysteines are annotated as on the cell surface in prior cysteinome datasets generated from whole cell proteomes^64^ (**Figure 1D**). Further showcasing the value of this dual capture method for identifying tough-to-detect residues, when compared with our CysDB database^63^, which aggregated a total of 62,888 cysteines in nine high coverage cysteine chemoproteomic datasets, 708 cysteines in proteins annotated as cell surface or glycosylated identified by Cys-Surf have not been previously identified (**Figure 1E**).

### Cys-Surf captures ligandable cell surface cysteines

Motivated by the robust capture of cell surface cysteines, we next set out to enable quantitative measures of subcellular cysteine ligandability by incorporating isotopic labeling into the Cys-Surf workflow. We envisioned that our previously reported isotopically labeled light- and heavy-isopropyl iodoacetamide alkyne (IPIAA-L **4** and IPIAA-H **5**), which had enabled high coverage bulk redox proteomics^70^, would prove compatible with competitive electrophilic small molecule screening. In a manner highly analogous to the widely utilized competitive chemoproteomic profiling platform - isotopic Tandem Orthogonal Proteolysis–Activity-Based Protein Profiling (isoTOP-ABPP)^66,82^, we treated samples in a pairwise manner with either scout fragment 3,5-bis(trifluoromethyl)aniline chloroacetamide **KB03** or vehicle (DMSO) followed by either light- or heavy-IPIAA (**Figure 2A**). After CSC and peptide level cysteine enrichment, FragPipe search and IonQuant^83^ was utilized to calculate the competition ration (ratio (R) of IPIAA-H over IPIAA-L).

**Figure 2.**
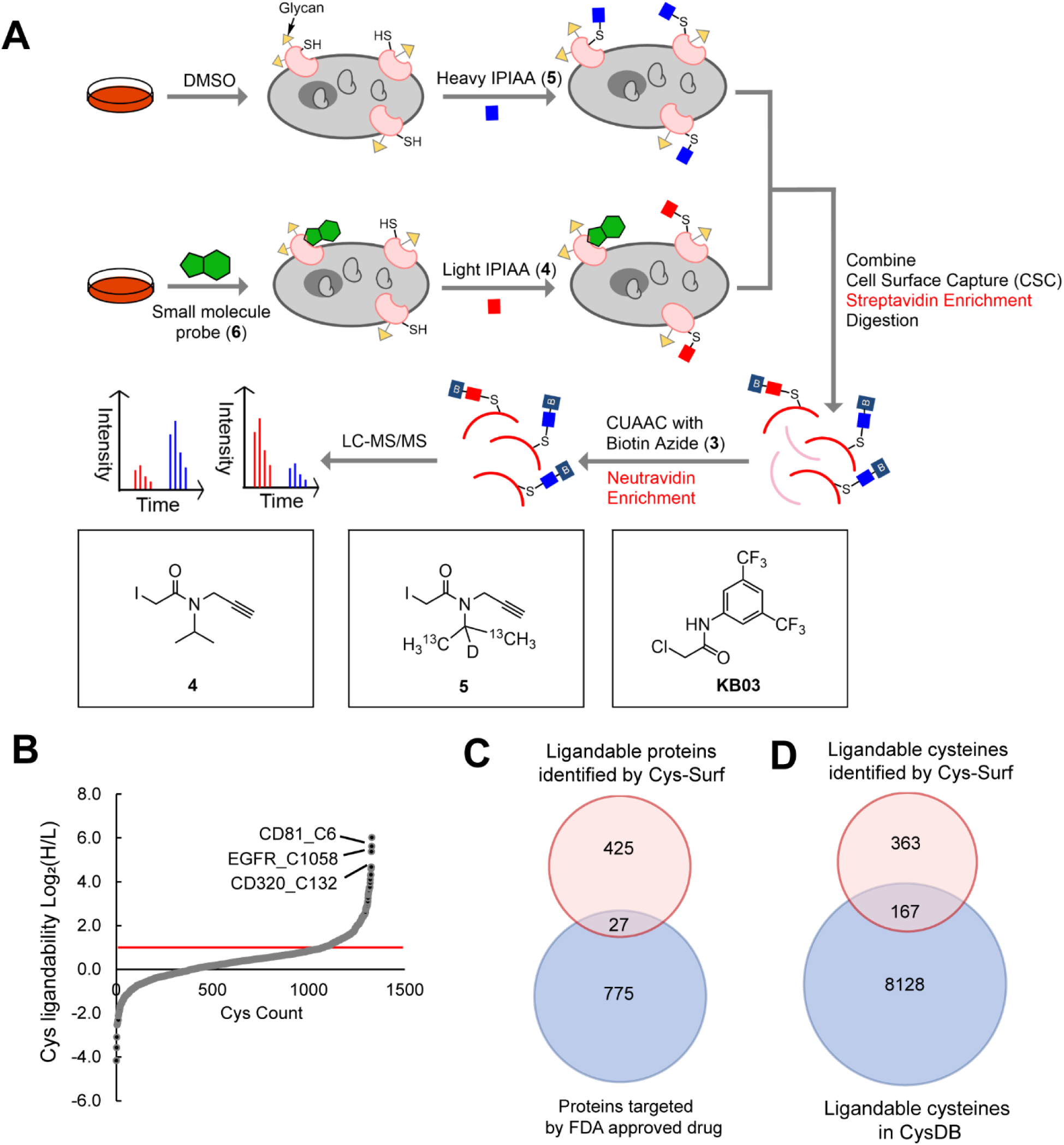
Application of Cys-Surf to identify ligandable cell surface cysteines. **A)** The schematic workflow to identify ligandable cell surface cysteines with Cys-Surf. **B)** Ligandability of cysteines quantified with Cys-Surf. **C)** Ligandable proteins identified by Cys-Surf compared to FDA approved drug targets. **D)** Ligandable cysteines identified by Cys-Surf compared to those in CysDB. Compound treatment was 50 μM for 1 h at 37 °C. MS experiments were conducted in 3 replicates in Jurkats cells. All MS data can be found in **Table S2**.

Out of the 2,194 cysteines quantified, 530 cysteines belonging to 452 proteins were found in peptides that showed elevated MS1 ratios (Log_2_(H/L) > 1), indicating covalent labeling by **KB03** (**Figure 2B**). Out of the 530 ligandable cysteines, 211 cysteines are annotated as from proteins on cell surface or glycosylated. Cys-Surf identified unique ligandable cell surface proteins that differ from those already targeted by FDA approved drugs (**Figure 2C**). More than 350 ligandable cell surface cysteines identified by our study are unique and have not been previously identified in CysDB^63^ (**Figure 3D**). While we do not identify C797 in EGFR—this is the cysteine labeled by targeted tyrosine kinase (TKIs) inhibitors (e.g. afatinib)^61^—we do observe an elevated ratio (Log_2_(H/L) = 2.74) for C1058 in EGFR. This cysteine has been previously identified as a palmitoylation site located in a disordered loop, which has been implicated in the palmitoylation-dependent regulation of EGFR activity^84,85^. Gene Ontology (GO)-analysis^86^ of proteins that harbor liganded cysteines demonstrated an enrichment of proteins involved in processes including integrin-mediated signaling pathway, membrane organization and biosynthesis (**Figure S4**). We expect that many of these ligandable cysteines, including C6 of CD81^87^ and C132 of CD320^88^, represent intriguing potential targets for immunomodulation^34^.

**Figure 3.**
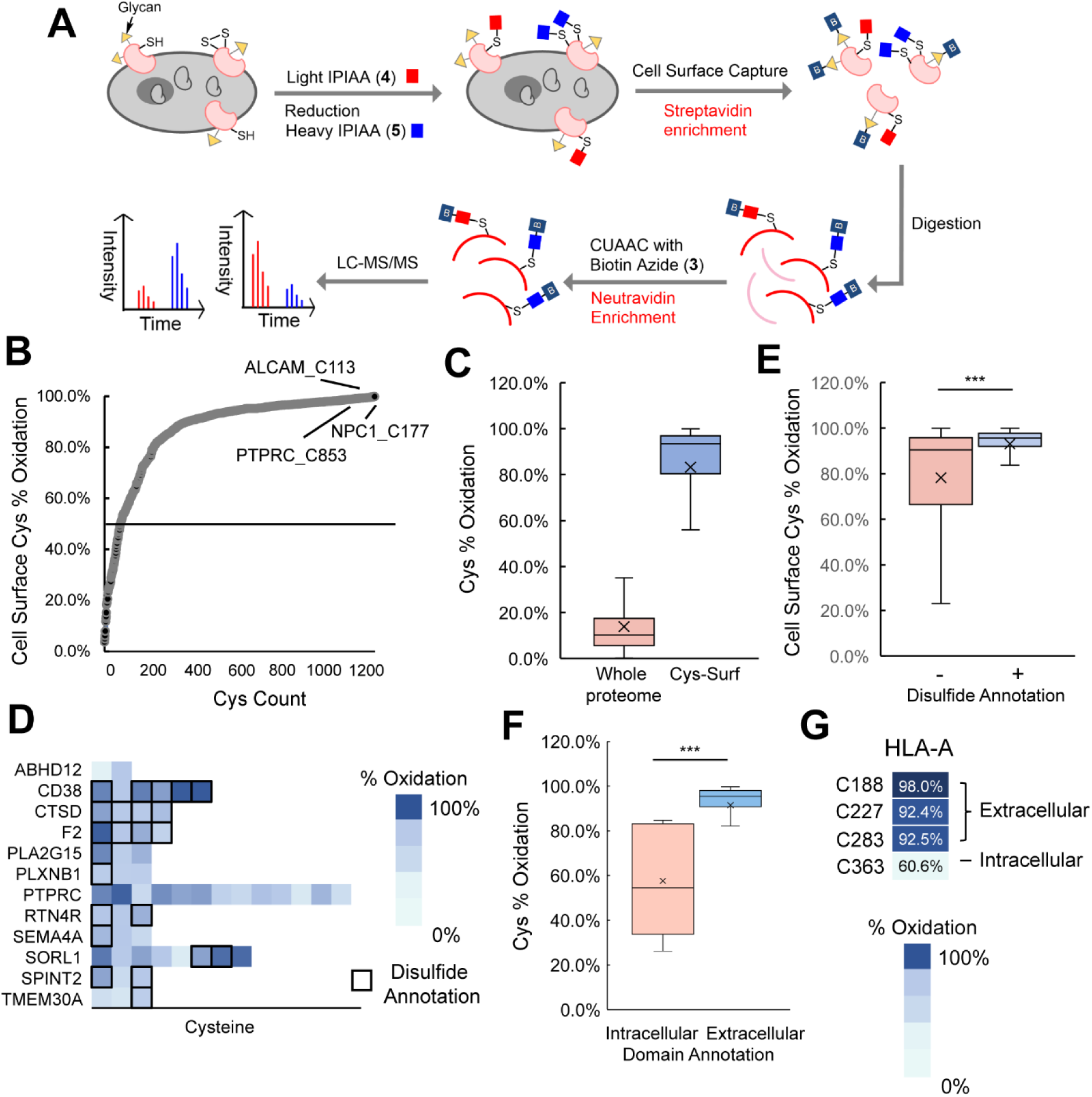
Application of Cys-Surf to quantify oxidation states of cell surface cysteines. **A)** The schematic workflow of quantification of oxidation states of cell surface cysteines with Cys-Surf. Reduced cysteines were first labeled using our custom isotopically light isopropyl iodoacetamide alkyne (IPIAA-L) probe. Subsequently, the samples were subjected to reduction and labeled using our isotopically differentiated heavy (IPIAA-H) probe. After CSC and peptide level cysteine enrichment, Fragpipe IonQuant reported the ratio (R) of IPIAA-H over IPIAA-L labeled cysteine peptides, and **B)** Percentage oxidation of cysteines quantified with Cys-Surf calculated using the formula (R/(1+R))*100 as reported previously^70,73^. Average percentage oxidation of 4 biological replicates is reported **C)** Percentage oxidation of cysteines quantified with Cys-Surf compared with that quantified with whole proteome. **D)** Percentage oxidation of representative cysteines quantified with Cys-Surf. **E)** Percentage oxidation of cysteines quantified with Cys-Surf with or without disulfide annotation. **F)** Percentage oxidation of cysteines quantified with Cys-Surf with domain analysis of extracellular or intracellular. **G)** Percentage oxidation of 4 cysteines in HLA-A quantified with Cys-Surf. For panel **E** and **F**, statistical significance was calculated with unpaired student’s t-tests, *** *p*<0.001. For panel **C**, **E**, and **F**, the mean is represented by “x”. MS experiments were conducted in 4 biological replicates in Jurkats cells. All MS data can be found in **Table S3**.

Among the integrin-related ligandable cysteines, C363 from HLA-A (HLA class I histocompatibility antigen; A-3 alpha chain) was identified as labeled by **KB03** (Log_2_(H/L) = 1.26). Inspection of the AlphaFold^89,90^ structure of HLA-A (AF-P04439-F1) revealed that C363 is located in a putatively unstructured c-terminal portion of the HLA protein (**Figure S5A**). Given the potential opportunities for targeting MHC complexes for immunomodulatory applications^91,92^, we opted to further assess the ligandability of this cysteine through a gel-based activity-based protein profiling (ABPP) assay^93^. We established an immunoprecipitation-based ABPP assay in which cells were first treated in situ for 1 h at 50 μM compound with one of three compounds, chloroacetamide (**KB03**), acrylamide (**KB14**) and propynamide (**6**). After lysis, lysates were labeled with IA-rhodamine and HLA-A protein was enriched by immunoprecipitation, which revealed that **6** labeled HLA-A, as indicated by the near-complete blockade of IA-Rho-labeling; more modest partial competition was observed of **KB03** and **KB14** (**Figure S5B**). Taken together, these findings support that Cys-Surf can faithfully identify ligandable cysteines within membrane-associated proteins.

### Quantitative MS analysis of oxidation states of cysteines on the cell surface with Cys-Surf

As the HLA-A Cys363 cysteine had been previously reported as redox active^94,95^, we next asked whether Cys-Surf would extend to quantification of site-specific cysteine oxidation. Building upon our previously reported SP3-Rox^70^ and Cys-LOx^73^ redox proteomics platforms, we modified our Cys-Surf platform to capture the cell surface redox proteome (**Figure 3A**). Similar to many redox proteomic methods^68,69,96,97^, we calculated the cysteine percentage oxidation state based on the relative cysteine labeling by our isotopically enriched IPIAA reagents, with IPIAA-L labeling reduced cysteines and IPIAA-H labeling oxidized cysteines following global reduction with DTT. In support of the robustness of our method, we also generated datasets in which the cysteine capping reagents were swapped, with IPIAA-H capping reduced cysteines and IPIAA-L capping the oxidatively modified cysteines after reduction. A high concordance (R^2 = 0.8942, n = 1,538 unique cysteines identified in both datasets) was revealed for the precursor ion intensity ratios calculated by FragPipe IonQuant^83^ (**Figure S6**).

Demonstrating the uniquely oxidizing environment of the cell surface cysteinome, 1,173 out of the 1,246 cysteines (89% of cysteines) with cell surface annotations quantified from 489 proteins were found to have elevated ratios (log_2_(H/L) >1), corresponding to >50% percent oxidation (**Figure 3B**). In aggregate, cysteines in cell surface proteins showed an average oxidation state of 83%; this high degree of overall oxidation is in stark contrast to the 10% mean cysteine oxidation state previously reported for whole cell proteome^70,72,82,98–101^, which supports Cys-Surf’s capacity to assay the subcellular oxidation state of cysteines in the plasma membrane (**Figure 3C**).

Given the markedly oxidized nature of most cysteines identified, we next asked whether these residues were known to be involved in formation of disulfide bonds. We find that 495 of the identified cysteines from 192 total proteins, were annotated as disulfide sites by UniProtKB. This coverage (40% of residues identified) represents a marked enrichment for cysteines involved in disulfides when compared to bulk proteome analysis—in CysDB 17% (n = 1,077) of identified cysteines are annotated as involved in disulfides. Exemplary cysteines involved in disulfide bonds include Cys177 of NPC intracellular cholesterol transporter 1 (NPC1), Cys296 of Cluster of Differentiation (CD) 38 antigen, and Cys113 of CD166 antigen ALCAM (**Figure 3B, 3D**).

Looking beyond these anecdotal examples, we find that the cysteines with annotated disulfide involvement showed a 93% mean oxidation, which reflects increased oxidation compared with all identified cell surface cysteines and to those lacking disulfide annotations (83% and 78% mean oxidation, respectively; **Figure 3E**). Notably, 453 additional cysteines, which lacked UniProtKB disulfide annotations, were also detected with >90% percent oxidization (**Table S3**). We expect that a number of these sites are likely cysteines involved in heretofore unannotated disulfides.

Given the predominance of highly oxidized cysteines in our dataset, we next asked whether Cys-Surf would have sufficient sensitivity to identify domain specific differences in cysteine oxidation. Using UniProtKB topological domain feature annotations, we identified that our dataset contained 175 cysteines located within annotated intracellular protein domains and 589 cysteines in annotated extracellular domains. Consistent with a high degree of oxidation of cysteines in the plasma membrane, we observe mean oxidation states of 92% for extracellular and versus 58% for intracellular domain cysteines (**Figure 3F**). For the aforementioned ligandable cysteine Cys363 in HLA-A, which is located in an annotated intracellular topological domain, we observe substantially reduced oxidation 61% when compared with the three HLA-A extracellular cysteines (Cys188, Cys227, and Cys283; **Figure 3G**).

### Cys-Surf identifies adaptive immune cell state-dependent changes to cysteine oxidation

During activation, T cells are exposed to reactive oxygen species (ROS) and reactive nitrogen species (RNS), derived from both intracellular mitochondrial ROS production and from extracellular neighboring phagocytes. Intriguingly, this oxidative environment has been connected with both T cell hypo-responsiveness as well as T cell activation, including increased IL2 and NFAT expression^102–104^. These seemingly antithetical activities of ROS as both pro- and anti-proliferative have been ascribed to both dose-dependent activity (low ROS is pro-proliferative and higher ROS-negatively impacting growth) and compartmentalized redox signaling. Intriguingly, while bulk proteome redox analysis revealed increased cysteine oxidation during T cell activation for a number of immunomodulatory targets^70,105,106^, an increase in cell surface cysteine free thiols is strongly associated with immunological stimuli^56,57,107–110^.

Inspired by these intriguing observations, we deployed Cys-Surf to pinpoint T cell activation-dependent changes to plasma membrane-associated cysteine oxidation state. Healthy donor T cells (bulk CD4+ and CD8+) were subjected to the Cys-Surf workflow (**Figure 4A**) pre- and post-stimulation (anti-CD3 and anti-CD28). In aggregate, the oxidation state of 2,402 cysteines on 1,190 proteins were quantified, in which 1,466 cysteines from 638 proteins are annotated as on the cell surface or are glycosylated (**Table S4**). Consistent with the aforementioned reports of activation-dependent increases in plasma membrane free thiol content, T cell stimulation afforded a modest yet significant decrease in the measured mean percentage cysteine oxidation state (**Figure 4B**). Out of 821 cysteines from 471 proteins quantified in both unstimulated and stimulated T cell proteomes, 100 cysteines were significantly reduced (difference < -1, p-value < 0.5) upon T cell stimulation (**Figure 4C**). GO-analysis^86^ of genes that harbor significantly reduced cysteines revealed an enrichment of proteins involved in processes including regulation of T cell tolerance induction, lymphocyte differentiation and integrin-mediated signaling, which are all closely related to T cell activation (**Figure 4D**). Exemplary reduced cell surface cysteines involved in adaptive immune response include a number of disulfide-bonded cysteines (**Figure 4E, S7**), including C41 from the T cell surface glycoprotein (CD4), which functions as a coreceptor for MHC class II molecule:peptide complex^57,111^, C70, C83 and C170 from the T cell differentiation antigen (CD6)^112,113^, and C385, C472 and C551 from the dipeptidyl peptidase 4 (DPP4), which is a known positive regulator of T cell activation^114–117^. For the receptor-type tyrosine-protein phosphatase C (PTPRC or CD45), which is a DPP4 interactor that also acts as a positive regulator of T cell coactivation, active site C853 that forms a phosphocysteine intermediate^118^ was markedly reduced. Notably, the disulfide-rich extracellular domains of CD45 have been implicated in increased domain rigidity and formation of heterodimers^119,120^. For CD4^121,122^ (Cys41) and Tapasin (TAPBP)^123^ (Cys91), the reduced disulfides are potential – RHStaple bonds—these are allosteric disulfides that link adjacent strands in the same β sheet^124,125^, which aligns with the comparatively labile nature of these bonds^126,127^. Exemplifying the value of subcellular redox measurements, for C551 of DPP4 and C170 of CD3, which were identified by both our previous bulk proteomic^70^- and our cys-Surf analysis of T cell activation, we observe markedly divergent state-dependent changes to thiol oxidation (**Figure 4E**).

**Figure 4.**
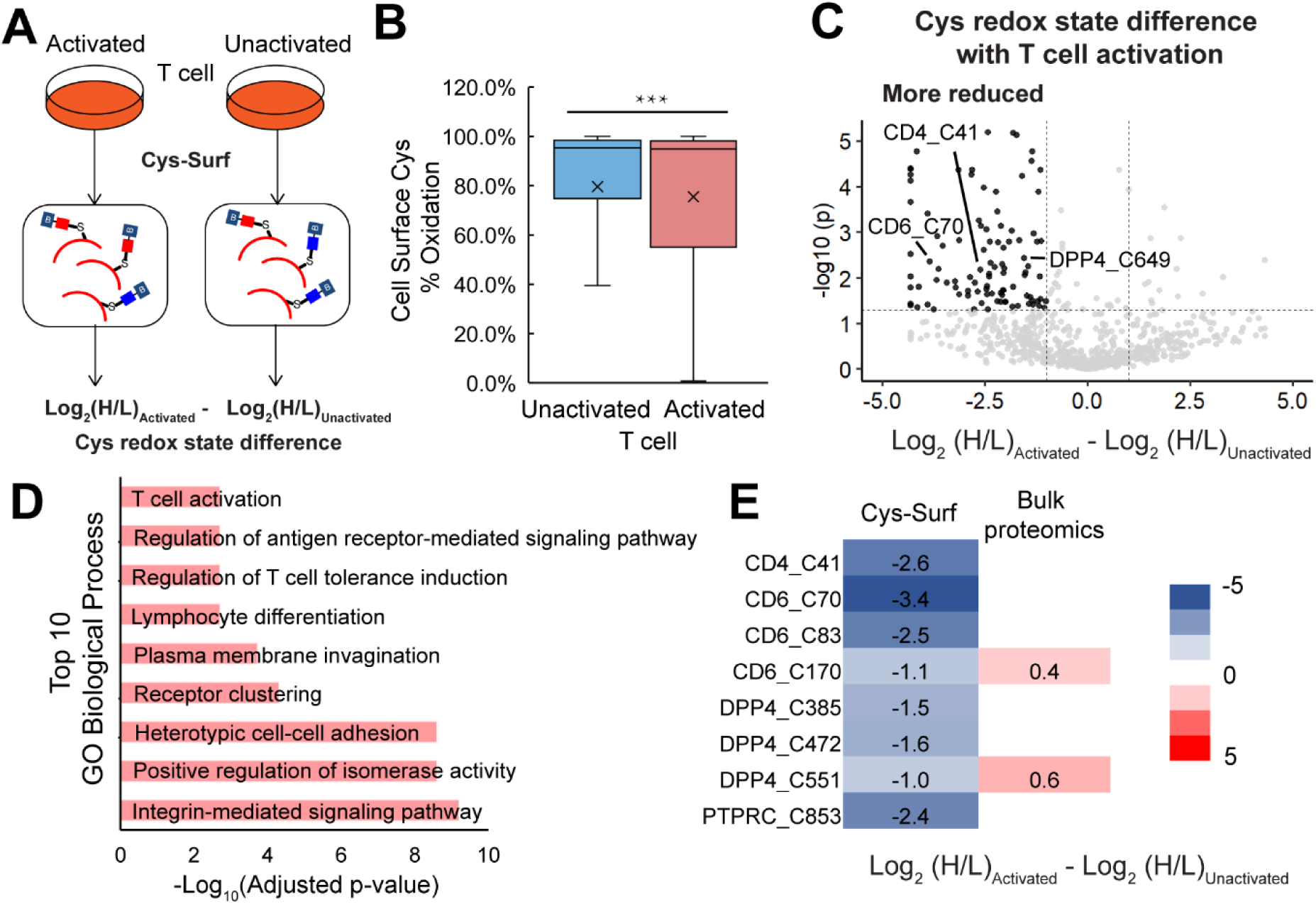
Application of Cys-Surf to quantify redox changes of cell surface cysteines during T cell activation. **A)** The schematic workflow to identify redox sensitive cell surface cysteines during T cell activation with Cys-Surf. **B)** Redox states of cell surface cysteines quantified with Cys-Surf in naive and activated T cells. The mean is represented by “x”. Statistical significance was calculated with unpaired student’s t-tests, *** *p*<0.001. **C)** Difference of redox states for cysteines quantified with Cys-Surf during T cell activation. **D)** Top 10 GO biological processes enriched with proteins harboring reduced cell surface cysteines during T cell activation. **E)** Difference of redox states of representative cysteines quantified with Cys-Surf or bulk proteomics in unactivated or activated T cells. MS experiments were conducted in 3 replicates in human T cells. All MS data can be found in **Table S4**.

### Cys-Surf identifies cysteines sensitive to exogenous reductants and oxidants

Inspired by the seeming ubiquity of labile disulfides, we opted to broaden the scope of redox sensitive cell surface cysteines by pairing Cys-Surf with cell treatments with exogenous oxidoreductants. For oxidants, we chose to the exemplary oxidizing agents H_2_O_2_ and S-Nitrosoglutathione (GSNO), with the goal of pinpointing cell surface cysteines sensitive to ROS and RNS-mediated cysteine oxidative modifications. For reductants, we selected both the pan-cysteine reactive Tris (2-carboxyethyl) phosphine (TCEP) reducing agents as well as N-acetylcysteine (NAC), which is a clinically administered reductant that is also widely taken as a dietary supplement^128,129^.

We subjected Jurkat cells to in-gel fluorescence analysis using the cell-impermeable cysteine-reactive Alexa Fluor™ 594 C_5_ Maleimide (Maleimide-Alexa594)^130,131^ to visualize treatment-dependent changes in cell surface cysteine reactivity (**Figure 5A**, top). For the H_2_O_2_- and GSNO-treated cells, we observed a modest decrease in the reactivity of free thiols towards Maleimide-Alexa594 (**Figure S8A**). In contrast, lysates derived from TCEP- and NAC-treated cells, exhibited increased Maleimide-Alexa594 labeling, consistent with reduction-dependent increased availability of free thiols. When compared to NAC, samples treated with TCEP exhibited a more substantial increase in Maleimide-Alexa594 reactivity, consistent with TCEP’s greater reduction potential (**Figure 5B**).

**Figure 5.**
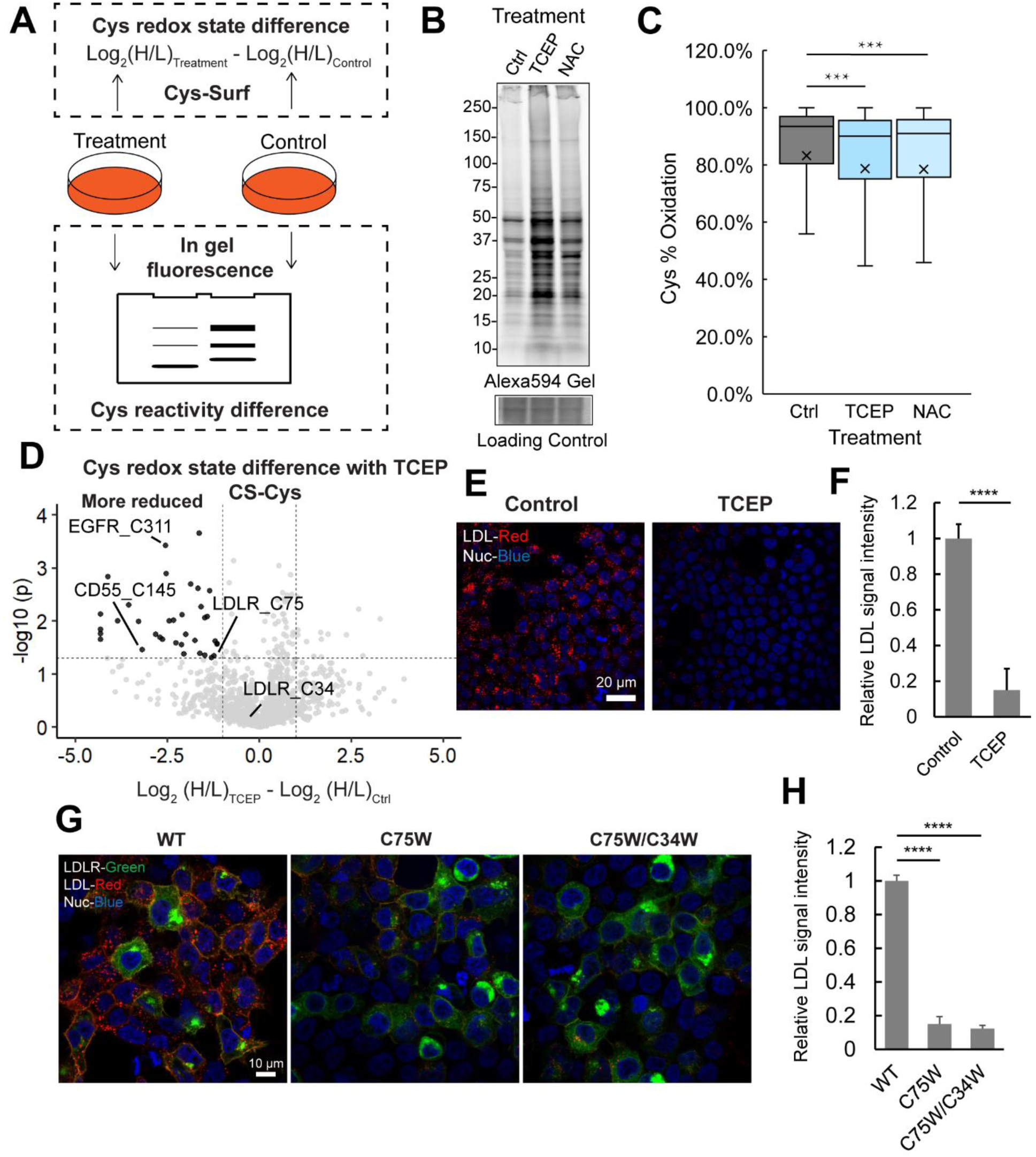
Application of Cys-Surf to identify redox sensitive cell surface cysteines. **A)** The schematic workflow to identify redox sensitive cell surface cysteines with Cys-Surf and in-gel fluorescence. **B)** Gel of cell surface cysteines labeled with Maleimide-Alexa594 treated with DMSO as control, tris(2-carboxyethyl)phosphine (TCEP) or N-acetyl cysteine (NAC). **C)** Redox states of cell surface cysteines quantified with Cys-Surf treated with DMSO, TCEP or NAC. The mean is represented by “x”. **D)** Difference of redox states for cysteines quantified with Cys-Surf with or without TCEP treatment. **E)** LDL uptake (signals in red) in HEK293T cells with or without treatment of TCEP. **F)** Quantification of relative LDL uptake signal intensities in HEK293T cells with or without treatment of TCEP. **G)** LDL uptake (signals in red) in HEK293T cells with expression of LDLR^WT^, LDLR^C75W^ and LDLR^C75W/C34W^ (signals in green). **H)** Quantification of relative LDL uptake signal intensities in HEK293T cells with expression of LDLR^WT^, LDLR^C75W^ and LDLR^C75W/C34W^ . TCEP treatment was 5 mM for 20 mins at 37 °C. NAC treatment was 10 mM for 1h at 37 °C. For panel **C**, **F**, and **H**, statistical significance was calculated with unpaired student’s t-tests, *** *p*<0.001, **** *p*<0.0005. For panel **F** and **H**, data is represented as mean ± stdev. MS experiments were conducted in 3 replicates in Jurkats cells. All MS data can be found in **Table S5**.

As our gel-based analyses confirmed treatment-dependent changes to cysteine oxidation, we next extended these analyses to Cys-Surf characterization of the oxidation- and reduction sensitive cell surface proteomes. Following the workflow shown in **Figure 5A** (bottom), we identified treatment-dependent changes to cysteine oxidation state based on the difference in MS1 intensity ratios comparing additive- and vehicle-treated samples. Consistent with the comparatively small decrease in Maleimide-Alexa594 labeling observed for oxidant treated samples (**Figure S8A**), Cys-Surf analysis of H_2_O_2_- and GSNO-treated cells revealed no significant treatment-dependent changes in the aggregated mean percentage oxidation of the cysteines identified (**Figure S8B**). These findings are consistent with our aforementioned observation (**Figure 2B, 2C**) that, at basal state, most cell surface cysteines are already highly (mean >80%) oxidized and thus insensitive to additional oxidant. This observed insensitivity is in stark contrast to that observed for whole proteome^70^, where we found that >51.0% of cysteines were significantly oxidized with GSNO. Out of 1,310 cysteines identified in both samples treated with vehicle and H_2_O_2_, 16 cysteines showed significant ratio changes (difference >1, p-value <0.05) sensitive to H_2_O_2_, including C590 of integrin beta-2 ITGB2, which has been reported as ligandable in human T cells^105^ (**Figure S8C**). Out of 305 cell surface cysteines identified in both samples with control and GSNO treatment, 3 cysteines showed significant GSNO induced oxidation (difference >1, p-value <0.05), including C109 of platelet endothelial cell adhesion molecule PECAM1, a protein that is required for leukocyte transendothelial migration (TEM) under most inflammatory conditions^132,133^ (**Figure S8D**).

Guided by the observed comparative insensitivity to exogenous oxidations and high degree of baseline oxidation state of cell surface cysteines, we next asked whether a more substantial fraction of the cell surface cysteinome would be sensitive to exogenous reductants TCEP and NAC. Consistent our in-gel fluorescence analysis (**Figure 5B**), Cys-Surf analysis revealed a modest yet significant decrease in the measured bulk mean cysteine oxidation state after TCEP and NAC treatment, decreasing from 93.4% to 90.0% and 91.0%, respectively (**Figure 5C**). Out of the 916 cysteines from 618 proteins identified in both samples with control and NAC treatment, 15 cysteines showed significant NAC induced reduction (difference <-1, p-value <0.05), including C173 of signaling threshold-regulating transmembrane adapter 1 SIT1, which has been reported as immune-enriched and hyperreactive in activated human T cells^105^ (**Figure S9**). Out of the 1,189 cysteines from 698 proteins identified in both samples treated with control and TCEP, 39 cysteines were sensitive to TCEP-induced to reduction (difference <-1, p-value <0.05), of which 32 has cell surface annotation. We initially were surprised by the low overlap between the NAC and TCEP datasets, with 534 shared total cysteines. Together with the difference in reducing potentials for TCEP and NAC, we expect that the comparatively modest coverage (1,573 total cysteines) together with the stochastic nature of data dependent acquisition (DDA) likely rationalizes the modest overlap between datasets.

Several intriguing reduction labile cysteines were identified only in the TCEP datasets. Annotated disulfides in multiple CD antigens were identified as reducible, including C145 of CD55, which has been identified as significantly reduced during T cell activation in our previous bulk proteome dataset^70^ (**Figure 5D, S10**). C311 of the epidermal growth factor receptor kinase (EGFR) was found to be markedly sensitive to TCEP (difference = -2.55). This cysteine is localized to a disulfide bridge (C311-C326) in a cysteine-rich region of EGFR that is required for kinase activation^134^. Notably, disulfide breaking mutations at C326 were recently reported to be gain-of-function for Lhermitte-Duclos Disease^135^.

Similarly, a large number of mutations at disulfide bonded extracellular cysteines in the low-density lipoprotein receptor (LDLR) have been implicated in Familial Hypercholesterolemia (FH), one of the most common genetic disorders^136,137^. In our TCEP dataset, we identified two cysteines from LDLR, highly reduced C75 (Difference = -1.5) and comparatively reduction insensitive C34 (Difference = -0.3). While variants at these two cysteines have been reported in ClinVar^138^, with clinical significance as likely pathogenic and associations with familial hypercholesteremia, the impact of these variants on LDL uptake is, to our knowledge, unknown. Notably, mutation at the proximal C46 has been demonstrated to reduce LDL uptake^139^.

Using fluorescently labeled LDL with GFP tagged on the C-term of LDL, we next assayed the impact of TCEP and C75 mutations on LDL uptake and LDLR activity. TCEP treatment attenuated both basal HEK-293T cellular LDL uptake and LDLR-dependent uptake with heterologously overexpressed LDLR (**Figure 5E, 5F, S11**). Guided by these findings, we then generated point mutations in our LDLR-GFP construct, including at C75 (LDLR^C75W^) and the double mutant (LDLR^C75W/C34W^), with the latter chosen to assess whether C75 mutations would synergize with the reduction-insensitive C34 residue.

HEK293T cells overexpressing the LDLR^C75W^ mutant showed markedly attenuated LDL uptake. In contrast to the substantial plasma membrane localization observed for the wild-type LDLR construct, we observed increased intracellularly localized LDLR^C75W^, consistent with mutation-induced changes to protein translocation to the plasma membrane. The double mutant (LDLR^C75W/C34W^) afforded a similar decrease in LDL uptake and intracellular protein retention (**Figure 5G, 5H, S12**). Taken together, our findings support the pairing of Cys-Surf with exogenous reductant treatments to pinpoint functional and reduction-labile disulfides. They also point towards future opportunities for therapeutic targeting of this comparatively underexplored portion of the cysteinome.

## Discussion

By establishing the cell <u>Surf</u>ace <u>Cys</u>teine enrichment (Cys-Surf) platform, here we achieved unprecedented local cysteinome analysis, including the identification of ligandable and redox sensitive membrane cysteines. Cys-Surf features a unique two-step biotinylation method: after in situ IAA capping to protect cysteines from oxidation, cell surface glycoproteins are then subjected to CSC on streptavidin followed by peptide-level click conjugation and neutravidin capture of cell surface-associated cysteine peptides. While our method draws inspiration from our recent pairing of TurboID proximity labeling with cysteine chemoproteomics^73^, to our knowledge such sequential rounds of enrichment for cell surface proteins is unprecedented. Application of Cys-Surf to primary immune cells and immortalized cell lines identified in aggregate 2,836 total cell surface cysteines from 1,046 proteins, including 211 ligandable cysteines and 153 redox sensitive cysteines. For more translational applications, we expect that our rich dataset of ligandable cell surface cysteines will be of particular interest, as demonstrated by the blockbuster status of several covalent kinase inhibitors that target cell surface-associated kinases (e.g. EGFR and HER2).

Enabled by our pairing of Cys-Surf with isotopically enriched cysteine capping reagents^70^, we stratified the cell surface redoxome. Consistent with prior studies, we find that the redox state of membrane associated cysteinome differs markedly from bulk proteomic analysis, both at basal state and for cells subjected to stimuli and exogenous oxidants and reductants. We find a mean oxidation state of ∼90% for the 1,246 cell surface cysteines for which quantitative measures of absolute oxidation were achieved. This striking oxidation contrasts with the previously reported 10% median cellular cysteine oxidation state, as measured by bulk redox proteomics^70,140,141^. Consistent with this marked oxidation and the highly oxidizing extracellular environment, we find that comparatively few cell surface cysteines are prone to oxidation, both with addition of exogenous oxidants and under cellular activation conditions. This latter finding is in contrast with the marked activation-induced oxidation observed for bulk proteomes^70^. These findings are consistent with previous reports of compartmentalization of redox-mediated signaling^142,143^ and highlight the value of subcellular measures of cysteine oxidation, as bulk proteome analysis will likely mask compartment-specific changes.

In contrast with the comparative insensitivity of the plasma membrane cysteine to oxidation, Cys-Surf revealed a prevalence of reduction labile cysteines, including many found in disulfide bonds within the cell surface proteome. Of the 53 total cysteines sensitive to the exogenous reductants TCEP and NAC, a number of noteworthy residues stood out, including those found in -RHStaples. In total we identified 21 disulfides sensitive to the exogenous reductants TCEP and NAC. Noteworthy examples of labile bonds included those found in multiple clusters of differentiation proteins implicated in cellular activation, a cysteine (C311) within an activation-associated region of EGFR, and a cysteine (C75) within the LDLR protein. While this cysteine (C75) of LDLR has been reported to be mutated in cases of Familial Hypercholesterolemia (FH), the functional impact of these mutations has not, to our knowledge, been probed previously. Providing compelling proof-of-concept evidence of the likely functional importance of these reduction-labile disulfide bonds, here, by combining mutational analysis and LDL uptake studies, we reveal that C75W mutations lead to retention of LDLR in the cytoplasm and decreased LDL uptake. While we chose the comparatively bulky tryptophan mutation to mimic small molecule labeling at this site, we expect that more conservative mutations should phenocopy our findings, as shown by other genetic variants that impact the cysteine-rich extracellular domain of LDLR^137,139^. Looking beyond LDLR, we anticipate that our work will help guide ongoing and future efforts to characterize the functional significance of labile disulfides, particularly in the context of receptor-mediated cell signaling.

Achieving a more comprehensive analysis of this intriguing fraction of the cysteinome will likely require improvements in coverage for future iterations of Cys-Surf. While the Cys-Surf coverage exceeded that of our TurboID-based platforms^73^, Cys-Surf cysteine peptide identification remained comparatively modest, with ∼1,000-2,000 cysteines captured per experiment. Thus, when compared with the typical >10,000 cysteines captured from bulk proteomic analysis, Cys-Surf is likely still substantially undersampling plasma membrane cysteines. Exemplifying this limitation, while we do identify cysteines from EGFR, we fail to capture C797, which is the cysteine modified by EGFR-directed covalent kinase inhibitors. Analysis of additional cell lines, including EGFR overexpressing cells, together with implementation of Cys-Surf with additional sequence specific proteases to improve coverage of cleave site poor transmembrane regions should afford enhanced coverage. Notably, despite this modest coverage, across the study, ∼30% (821/2836) of the cysteines identified by Cys-Surf were not found previously in CysDB, our database of high coverage cysteine chemoproteomic datasets.

Looking beyond the current study, we expect that Cys-Surf analysis should synergize with genetic^144^ and proteogenomic^145^ approaches aimed at pinpointing functional and therapeutically relevant cysteines, including those impacted by genetic variation. Given the seeming ubiquity of reduction-labile disulfide bonds identified by Cys-Surf, we expect that a subset of these cysteines should be susceptible to covalent inhibitor development efforts, including a subset of the >500 ligandable cysteines identified here. While covalent inhibition of labile-disulfide-bonds is precedented by molecules targeting oxidoreductase enzymes^146,147^, the scope and functional impact of targeting redox environment-dependent cysteine proteoforms remain to be realized. The accessibility of extracellular cysteines identified by Cys-Surf to biologic agents with poor cell penetrance, including peptides and antibodies, offers a unique opportunity for the future development of cysteine-directed biological agents, as well as proteoform-specific therapies.

## Acknowledgements

This study was supported by a Packard Fellowship (K.M.B.), DP2 OD030950-01 (K.M.B.), National Institutes of Health 1P01HL146358-01 (P.T., K.M.B.), TRDRP T31DT1800 (T.Y.), and NIGMS System and Integrative Biology 5T32GM008185-33 (L.M.B.). We thank all members of the Backus labs for helpful suggestions and the CFAR virology core 5P30 AI028697 for providing donor T cells.

## Author Contributions

T.Y. and K.M.B. conceived of the project. T.Y. and L.J. collected data. T.Y. and L.M.B. performed data analysis. L.M.B. wrote software. T.Y. contributed to the figures. T.Y. and K.M.B. wrote the manuscript with assistance from all authors.

## Methods

### Cell culture

Cell culture reagents including Dulbecco’s phosphate-buffered saline (DPBS), Roswell Park Memorial Institute (RPMI) media, Dulbecco’s Modified Eagle Medium (DMEM) media and penicillin/streptomycin (Pen/Strep) were purchased from Fisher Scientific. Fetal Bovine Serum (FBS) was purchased from Avantor Seradigm (lot # 214B17). All cell lines were obtained from ATCC, routinely tested for mycoplasma, and were maintained at a low passage number (< 20 passages). Jurkat (ATCC: TIB-152) cells were cultured in RPMI-1640 supplemented with 10% FBS and 1% antibiotics (Penn/Strep, 100 U/mL). HEK293T (ATCC: CRL-3216) cells were cultured in DMEM supplemented with 10% FBS and 1% antibiotics (Penn/Strep, 100 U/mL). Media was filtered (0.22 μm) prior to use. Cells were maintained in a humidified incubator at 37 °C with 5% CO2.

Blood from a healthy donor was obtained from UCLA/CFAR Virology Core (5P30 AI028697) after informed consent. After Trima filter isolation, peripheral blood mononuclear cells (PBMCs) were purified over Ficoll–Hypaque gradients (Sigma-Aldrich) and T cells were purified via negative selection with magnetic beads (EasySep Human T Cell Iso Kit, 17951, STEMCELL). The purified T cells were washed with sterile PBS. Unstimulated cells were harvested by centrifugation. The remaining cells were then resuspended in RPMI-1640 supplemented with FBS, penicillin, streptomycin and glutamine (2 million cells per ml) and 200,000 cells per well were seeded on non-treated tissue culture, 96-well transparent plates that had been coated with anti-CD3 (1:200, BioXcell) and anti-CD28 (1:500, Biolegend) in PBS (100 μl per well). After 72h, the cells were harvested by centrifugation (4,500 *g*, 5 min, 4 °C), washed 3 times with cold DPBS and prepared as described below.

### Proteomic sample preparation for Cys-Surf, protein level enrichment

After cells were treated as indicated, harvested by centrifugation (4,500 *g*, 5 min, 4 °C) and washed with DPBS for 3 times, they were resuspended in DPBS and labeled with 2 mM IAA for 1h at room temperature (RT). For cysteine oxidation quantification, cells were labeled with 2 mM L-IPIAA for 1h at RT, reduced with 1 mM DTT for 15 min at 37 °C, and labeled with 2 mM H-IPIAA for 1h at RT, with PBS wash between the IPIAA labeling. For cysteine ligandability quantification, cells were labeled with either compound or vehicle for 1h at RT, followed by treatment of 2 mM L-IPIAA or H-IPIAA for 1h at RT respectively, and then combined. Following cysteine capture, cells were oxidized with 1.6 mM NaIO_4_ in 1 mM NaAc (pH = 6.5) for 20 min at RT in dark. After washing with DPBS, cells were labeled with 1 mM aminooxy-biotin in PBS for 1h at 4 °C. Cells were then washed with DPBS, lysed in RIPA buffer (Fisher, Cat# AAJ62885AE) for 30 min at 4 °C, and clarified by centrifuging (21,000 *g*, 10 min, 4 °C). Protein concentrations were determined using a BCA protein assay kit (Thermo Fisher, Cat# 23227) and the lysate were normalized to 500 μL of 1.5 mg/mL. 50 μL Sera-Mag SpeedBeads Carboxyl Magnetic Beads, hydrophobic (GE Healthcare, 65152105050250) and 50 μL Sera-Mag SpeedBeads Carboxyl Magnetic Beads, hydrophilic (GE Healthcare, 45152105050250) were mixed and washed with water three times. The bead slurries were then transferred to the lysate, incubated for 5 min at RT with shaking (1000 rpm). 1 mL EtOH was added to each sample and the mixtures were incubated for 10 min at RT with shaking (1000 rpm). The beads were then washed (3 × 1 mL 80% EtOH) with a magnetic rack. Proteins were eluted from SP3 beads with 500 μL of 0.2% SDS in PBS for 30 min at 37 °C with shaking (1000 rpm). 50 μL Pierce streptavidin agarose beads were washed with 0.2% SDS/PBS and incubated with the eluted lysates for 2 h at RT. The proteins bound to beads were washed once with 1 mL 0.2% SDS/PBS, 3 times with 1 mL PBS, and 3 times with 1 mL H_2_O. The beads were resuspended in 200 μL 6 M urea, either reduced with 1 mM DTT for 15 min at 65 °C, and labeled with 2 mM IAA for 1h at RT for Cys-CS identification, or reduced with 10 mM DTT for 15 min at 65 °C, and labeled with 20 mM IA for 30 min at 37 °C for Cys-CS oxidation and ligandability quantification. Then, beads were washed with PBS and resuspended in 200 μL 2 M urea. 3 μL of 1 mg/mL trypsin solution (Washington) was added. Proteins were digested off the bead overnight at 37 °C with shaking.

### Proteomic sample preparation for Cys-Surf, peptide level enrichment

After digestion, CuAAC was performed with biotin-azide (4 μL of 200 mM stock in DMSO, final concentration = 4 mM), TCEP (4 μL of fresh 50 mM stock in water, final concentration = 1 mM), TBTA (12 μL of 1.7 mM stock in DMSO/*t*-butanol 1:4, final concentration = 100 μM), and CuSO4 (4 μL of 50 mM stock in water, final concentration = 1 mM) for 1h at RT. 20 μL Sera-Mag SpeedBeads Carboxyl Magnetic Beads, hydrophobic and 20 μL Sera-Mag SpeedBeads Carboxyl Magnetic Beads, hydrophilic were mixed and washed with water three times. The bead slurries were then transferred to the CuAAC samples, incubated for 5 min at RT with shaking (1000 rpm). Approximately 4 mL acetonitrile (> 95% of the final volume) was added to each sample and the mixtures were incubated for 10 min at RT with shaking (1000 rpm). The beads were then washed (3 × 1 mL acetonitrile) with a magnetic rack. Peptides were eluted from SP3 beads with 100 μL of 2% DMSO in MB water for 30 min at 37 °C with shaking (1000 rpm). The elution was repeated again with 100 μL of 2% DMSO in MB water. For each sample, 50 μL of NeutrAvidin Agarose resin slurry (Pierce, 29200) was washed three times in 10 mL IAP buffer (50 mM MOPS pH 7.2, 10 mM sodium phosphate, and 50 mM NaCl buffer) and then resuspended in 800 μL IAP buffer. Peptide solutions eluted from SP3 beads were then transferred to the NeutrAvidin Agarose resin suspension, and the samples were rotated for 2 h at RT. After incubation, the beads were pelleted by centrifugation (2,000 *g*, 1 min) and washed (3 × 1 mL PBS, 3 × 1 mL water). Bound peptides were eluted twice with 60 μL of 80% acetonitrile in MB water containing 0.1% FA. The first 10 min incubation at RT and the second one at 72 °C. The combined eluants were dried (SpeedVac), then reconstituted with 5% acetonitrile and 1% FA in MB water and analyzed by LC-MS/MS.

### Database Construction

Subcellular location annotations and glycosylation annotations from CellWhere Atlas (accessed 2208), Human Protein Atlas (HPA) version 21.1 and UniProtKB/Swiss-Prot (2208_release) were aggregated. Unique proteins were established using UniProt protein identifiers. Aggregated annotations were mined for specific keywords (ex. ‘Cell Surface’ or ‘Glycosylation’). Proteins containing these keywords are reported in **Table S1**.

### Gene Enrichment analysis

Enrichment of Kyoto Encyclopedia of Genes and Genomes (KEGG) 2021 and Gene Ontology (GO) Biological Processes 2021 gene set library terms were performed using the GSEApy package^148^. Proteins identified by our chemoproteomics studies were utilized as the background protein set. UniProtKB protein identifiers were mapped to Entrez gene symbols as input for Enrichr. P-values were computed from Fisher’s exact test to determine the significance of each enriched term. The negative log of these p-values was calculated using R.

### Cell treatment

H_2_O_2_: Cells were treated with 500 μM H_2_O_2_ for 1h at 37 °C. S-nitrosoglutathione (GSNO): Cells were treated with 500 μM GSNO for 1h at 37 °C. Tris(2-carboxyethyl)phosphine (TCEP): Cells were treated with 5 mM TCEP for 20 mins at 37 °C. N-acetylcysteine (NAC): Cells were treated with 10 mM NAC for 1h at 37 °C. 3,5-bis(trifluoromethyl)aniline chloroacetamide (**KB03**), acrylamide (**KB14**) and propynamide (**6**): Cells were treated with 50 μM compound for 1h at 37 °C.

### Gel and western blot

For streptavidin blot, lysates after Cell Surface Capture (CSC) were normalized to 2 mg/mL and separated on a 4-20% SDS-PAGE gel. Gels were transferred to nitrocellulose membrane (Bio-Rad) and blocked in 2% (w/v) BSA in TBS-T (Tris-buffered saline, 0.1% Tween 20) for 1h at RT. Membranes were incubated with a streptavidin-fluorophore conjugate overnight at 4 °C. Membranes were imaged on Bio-Rad ChemiDoc. For fluorescent gel, cells were treated as described and labeled with 5 μM IA-Rho or 5 μM Maleimide-Alexa594. After lysis, lysates were normalized to 2 mg/mL and separated on a 4-20% SDS-PAGE gel. Gels were imaged on Bio-Rad ChemiDoc. Reagents can be found in **Table S6**.

### Immunoprecipitation

Cell lysates were incubated with antibody overnight at 4 °C. 25 μL protein g beads were washed, added to the lysate and incubated at 4 °C for 2 h with rotation. After washing the beads with lysis buffer for 3 times, 25 μL of SDS loading dye was added to the sample and incubated at 95 °C for 5 min. Supernatant was collected for gel analysis after centrifugation at 12,000 *g* for 2 min.

### Construction of LDLR plasmids

Human LDLR gene was cloned from HepG2 cell cDNA and was then sequentially subcloned into pEGFP-N1 using the Gateway technology (Invitrogen). The LDLR mutations for the pEGFP-N1-LDLR construct were introduced by site-directed mutagenesis.

### LDL uptake assay and microscopy

HEK293T cells were plated in a standard 24-well plate, with each well containing a poly-d-lysine-coated glass coverslip (NC0672873, Fisher Scientific). Cells were transfected with LDLR-GFP (WT, C75W, and C75W/C34W) for 48 h with Fugene 6 (Promega). Cells were starved in 5% LPDS media (Millipore Sigma, S5519) for 12 h. Cells were then incubated with 10 µg/mL Dil-LDL (Kalen Biomedical, 770230-9) for 50 min. After Dil-LDL incubation, cells were washed three times with DPBS and fixed with 4% PFA for 20 min at room temperature. After fixation, cells were washed three times with DPBS and incubated with Hoechst 33342 (H3570, Fisher Scientific). Cells were mounted on slides with Prolong Diamond Antifade Mountant (P36961, ThermoFisher). Images were acquired using an Inverted Leica TCS-SP8 Confocal Microscope (CNSI) and analyzed by Image J.

### Liquid-chromatography tandem mass-spectrometry (LC-MS/MS) analysis

The samples were analyzed by liquid chromatography tandem mass spectrometry using a Thermo Scientific™ Orbitrap Eclipse™ Tribrid™ mass spectrometer. Peptides were fractionated online using a 18 cm long, 100 μM inner diameter (ID) fused silica capillary packed in-house with bulk C18 reversed phase resin (particle size, 1.9 μm; pore size, 100 Å; Dr. Maisch GmbH). The 70-minute water-acetonitrile gradient was delivered using a Thermo Scientific™ EASY-nLC™ 1200 system at different flow rates (Buffer A: water with 3% DMSO and 0.1% formic acid and Buffer B: 80% acetonitrile with 3% DMSO and 0.1% formic acid). The detailed gradient includes 0 – 5 min from 3 % to 10 % at 300 nL/min, 5 – 64 min from 10 % to 50 % at 220 nL/min, and 64 – 70 min from 50 % to 95 % at 250 nL/min buffer B in buffer A (**Table S7**). Data was collected with charge exclusion (1, 8, >8). Data was acquired using a Data-Dependent Acquisition (DDA) method consisting of a full MS1 scan (Resolution = 120,000) followed by sequential MS2 scans (Resolution = 15,000) to utilize the remainder of the 1 second cycle time. Precursor isolation window was set as 1.6 and normalized collision energy was set as 30%. The MS data have been deposited to the ProteomeXchange Consortium via the PRIDE partner repository^149,150^ with the dataset identifier PXD042403. File details can be found in **Table S8**.

### Protein, peptide, and cysteine identification

Raw data collected by LC-MS/MS were converted to mzML and searched with MSFragger (v3.3) and FragPipe (v19.0)^151^. The proteomic workflow and its collection of tools was set as default and PTMprophet was enabled^152^. Precursor and fragment mass tolerance was set as 20 ppm. Missed cleavages were allowed up to 1. Peptide length was set 7 - 50 and peptide mass range was set 500 - 5000. For Cys-Surf identification, cysteine residues were searched with differential modification C+463.2366. For Cys-Surf oxidation and ligandability quantification, MS1 labeling quant was enabled with Light set as C+463.2366 and Heavy set as C+467.2529. MS1 intensity ratio of heavy and light labeled cysteine peptides were reported with Ionquant (v1.8.9). Calibrated and deisotoped spectrum files produced by FragPipe were retained and reused for this analysis. The MS search data have been deposited to the ProteomeXchange Consortium via the PRIDE partner repository with the dataset identifier PXD042403. File details can be found in **Table S8**. Custom python scripts were implemented to compile labeled peptide datasets. Unique proteins, unique cysteines, and unique peptides were quantified for each dataset. Unique proteins were established based on UniProt protein IDs. Unique peptides were found based on sequences containing a modified cysteine residue. Unique cysteines were classified by an identifier consisting of a UniProt protein ID and the amino acid number of the modified cysteine (ProteinID_C#); residue numbers were found by aligning the peptide sequence to the corresponding UniProt protein sequence. When there are multiple cysteines in one peptide, all the modified cysteine residue numbers will be reported as ProteinID_C#_C#.

### Data analysis

For the cell surface annotation, our customized localization database was used to cross referenced with the proteins or cysteines identified. For isotopical quantification, the medium of heavy over light ratios for the same cysteine residue from cysteine peptides of different charges and miss cleavages in the same sample was calculated. Means of reported logged ratio values for each condition (+/- H_2_O_2_ or +/- GSNO or +/- TCEP or +/-NAC or activated/naive T cells) were calculated for all replicates per condition. Percentage oxidation for a cysteine was calculation based on heavy to light ratio via the following formula: (R/(1+R))*100, using unlogged ratios. When calculating oxidation difference, relative oxidation changes between two cellular conditions were reported by calculating the change of heavy to light ratios between treated and untreated samples.

### UniProtKB disulfide and domain analysis

Counts of how many identified proteins had UniProtKB annotations for disulfide bonds and topological domain annotations were calculated based on matches between the position of the identified residue and UniProtKB functional region. Further parsing of UniProtKB disulfide bond site annotations were extracted to obtain specific residues and amino acid numbers. Exact amino acid positions of UniProtKB cysteines involved in disulfide bonds or within regions of annotated domains were cross-referenced with our cysteine identifiers. Cysteines with ‘extracellular’ topological domain annotations were classified as cysteines in extracellular domains, while cysteines with ‘cytoplasmic’, ‘mitochondrial intermembrane’, ‘mitochondrial matrix’, and ‘perinuclear space’ were classified as cysteines in intracellular domains.

### Statistics

For box plots in **Figure 3C, 3E, 3F, 4B, 5C and S8B**, average of replicates was reported as indicated. Statistical significance was calculated with unpaired Student’s t-tests using R stats (v 3.6.2) if applicable. For bar plots in **Figure 1D, 5F, 5H, S1E, S2B and S3**. Error bars were calculated using standard deviation. Statistical significance was calculated with unpaired Student’s t-tests using R stats (v 3.6.2) if applicable. For volcano plots in **Figure 4C, 5D, S8C, S8D and S9A**, variances were calculated for each sample-condition pairing and a corresponding two-sample t-test was performed using R stats (v 3.6.2) to generate p-values. p-values were adjusted for multiple comparisons using Benjamini-Hochberg procedure.

## References

1. Li, J.; Han, S.; Li, H.; Udeshi, N. D.; Svinkina, T.; Mani, D. R.; Xu, C.; Guajardo, R.; Xie, Q.; Li, T.; Luginbuhl, D. J.; Wu, B.; McLaughlin, C. N.; Xie, A.; Kaewsapsak, P.; Quake, S. R.; Carr, S. A.; Ting, A. Y.; Luo, L. Cell-Surface Proteomic Profiling in the Fly Brain Uncovers Wiring Regulators. Cell 2020, 180 (2), 373–386.e15.

2. da Cunha, J. P. C.; Galante, P. A. F.; de Souza, J. E.; de Souza, R. F.; Carvalho, P. M.; Ohara, D. T.; Moura, R. P.; Oba-Shinja, S. M.; Marie, S. K. N.; Silva, W. A.; Perez, R. O.; Stransky, B.; Pieprzyk, M.; Moore, J.; Caballero, O.; Gama-Rodrigues, J.; Habr-Gama, A.; Kuo, W. P.; Simpson, A. J.; Camargo, A. A.; Old, L. J.; de Souza, S. J. Bioinformatics Construction of the Human Cell Surfaceome. Proc. Natl. Acad. Sci. 2009, 106 (39), 16752– 16757.

3. Shuster, S. A.; Li, J.; Chon, Ur.; Sinantha-Hu, M. C.; Luginbuhl, D. J.; Udeshi, N. D.; Carey, D. K.; Takeo, Y. H.; Xie, Q.; Xu, C.; Mani, D. R.; Han, S.; Ting, A. Y.; Carr, S. A.; Luo, L. In Situ Cell-Type-Specific Cell-Surface Proteomic Profiling in Mice. Neuron 2022, 110 (23), 3882–3896.e9.

4. Wojdyla, K.; Collier, A. J.; Fabian, C.; Nisi, P. S.; Biggins, L.; Oxley, D.; Rugg-Gunn, P. J. Cell-Surface Proteomics Identifies Differences in Signaling and Adhesion Protein Expression between Naive and Primed Human Pluripotent Stem Cells. Stem Cell Reports 2020, 14 (5), 972–988.

5. Gaultier, A.; Simon, G.; Niessen, S.; Dix, M.; Takimoto, S.; Cravatt, B. F.; Gonias, S. L. LDL Receptor-Related Protein 1 Regulates the Abundance of Diverse Cell-Signaling Proteins in the Plasma Membrane Proteome. J. Proteome Res. 2010, 9 (12), 6689–6695.

6. Schumann, J. Molecular Mechanism of Cellular Membranes for Signal Transduction. Membranes (Basel). 2022, 12 (8), 748.

7. Queiroz, R. M. L.; Charneau, S.; Bastos, I. M. D.; Santana, J. M.; Sousa, M. V.; Roepstorff, P.; Ricart, C. A. O. Cell Surface Proteome Analysis of Human-Hosted Trypanosoma Cruzi Life Stages. J. Proteome Res. 2014, 13 (8), 3530–3541.

8. Ravenhill, B. J.; Soday, L.; Houghton, J.; Antrobus, R.; Weekes, M. P. Comprehensive Cell Surface Proteomics Defines Markers of Classical, Intermediate and Non-Classical Monocytes. Sci. Rep. 2020, 10 (1), 4560.

9. Soday, L.; Potts, M.; Hunter, L. M.; Ravenhill, B. J.; Houghton, J. W.; Williamson, J. C.; Antrobus, R.; Wills, M. R.; Matheson, N. J.; Weekes, M. P. Comparative Cell Surface Proteomic Analysis of the Primary Human T Cell and Monocyte Responses to Type I Interferon. Front. Immunol. 2021. 10.3389/fimmu.2021.600056.

10. Schiess, R.; Mueller, L. N.; Schmidt, A.; Mueller, M.; Wollscheid, B.; Aebersold, R. Analysis of Cell Surface Proteome Changes via Label-Free, Quantitative Mass Spectrometry. Mol. Cell. Proteomics 2009, 8 (4), 624–638.

11. Karcini, A.; Lazar, I. M. The SKBR3 Cell-Membrane Proteome Reveals Telltales of Aberrant Cancer Cell Proliferation and Targets for Precision Medicine Applications. Sci. Rep. 2022, 12 (1), 10847.

12. Özlü, N.; Qureshi, M. H.; Toyoda, Y.; Renard, B. Y.; Mollaoglu, G.; Özkan, N. E.; Bulbul, S.; Poser, I.; Timm, W.; Hyman, A. A.; Mitchison, T. J.; Steen, J. A. Quantitative Comparison of a Human Cancer Cell Surface Proteome between Interphase and Mitosis. EMBO J. 2015, 34 (2), 251–265.

13. Affara, M.; Dunmore, B.; Savoie, C.; Imoto, S.; Tamada, Y.; Araki, H.; Charnock-Jones, D. S.; Miyano, S.; Print, C. Understanding Endothelial Cell Apoptosis: What Can the Transcriptome, Glycome and Proteome Reveal? Philos. Trans. R. Soc. B Biol. Sci. 2007, 362 (1484), 1469–1487.

14. Drews, J. Drug Discovery: A Historical Perspective. Science (80-.). 2000, 287 (5460), 1960–1964.

15. Overington, J. P.; Al-Lazikani, B.; Hopkins, A. L. How Many Drug Targets Are There? Nat. Rev. Drug Discov. 2006, 5 (12), 993–996.

16. Collier, A. J.; Panula, S. P.; Schell, J. P.; Chovanec, P.; Plaza Reyes, A.; Petropoulos, S.; Corcoran, A. E.; Walker, R.; Douagi, I.; Lanner, F.; Rugg-Gunn, P. J. Comprehensive Cell Surface Protein Profiling Identifies Specific Markers of Human Naive and Primed Pluripotent States. Cell Stem Cell 2017, 20 (6), 874–890.e7.

17. Gundry, R. L.; Boheler, K. R.; Van Eyk, J. E.; Wollscheid, B. A Novel Role for Proteomics in the Discovery of Cell-Surface Markers on Stem Cells: Scratching the Surface. PROTEOMICS – Clin. Appl. 2008, 2 (6), 892–903.

18. Kuhlmann, L.; Cummins, E.; Samudio, I.; Kislinger, T. Cell-Surface Proteomics for the Identification of Novel Therapeutic Targets in Cancer. Expert Rev. Proteomics 2018, 15 (3), 259–275.

19. McEnaney, P. J.; Parker, C. G.; Zhang, A. X.; Spiegel, D. A. Antibody-Recruiting Molecules: An Emerging Paradigm for Engaging Immune Function in Treating Human Disease. ACS Chem. Biol. 2012, 7 (7), 1139–1151.

20. Elschenbroich, S.; Kim, Y.; Medin, J. A.; Kislinger, T. Isolation of Cell Surface Proteins for Mass Spectrometry-Based Proteomics. Expert Review of Proteomics. 2010, pp 141–154.

21. Macher, B. A.; Yen, T.-Y. Proteins at Membrane Surfaces—a Review of Approaches. Mol. Biosyst. 2007, 3 (10), 705.

22. Wu, C. C.; Yates, J. R. The Application of Mass Spectrometry to Membrane Proteomics. Nat. Biotechnol. 2003, 21 (3), 262–267.

23. Vuckovic, D.; Dagley, L. F.; Purcell, A. W.; Emili, A. Membrane Proteomics by High Performance Liquid Chromatography-Tandem Mass Spectrometry: Analytical Approaches and Challenges. Proteomics. 2013, pp 404–423.

24. Elia, G. Biotinylation Reagents for the Study of Cell Surface Proteins. Proteomics 2008, 8 (19), 4012–4024.

25. Rybak, J.-N.; Ettorre, A.; Kaissling, B.; Giavazzi, R.; Neri, D.; Elia, G. In Vivo Protein Biotinylation for Identification of Organ-Specific Antigens Accessible from the Vasculature. Nat. Methods 2005, 2 (4), 291–298.

26. Li, Y.; Wang, Y.; Mao, J.; Yao, Y.; Wang, K.; Qiao, Q.; Fang, Z.; Ye, M. Sensitive Profiling of Cell Surface Proteome by Using an Optimized Biotinylation Method. J. Proteomics 2019, 196, 33–41.

27. Pischedda, F.; Szczurkowska, J.; Cirnaru, M. D.; Giesert, F.; Vezzoli, E.; Ueffing, M.; Sala, C.; Francolini, M.; Hauck, S. M.; Cancedda, L.; Piccoli, G. A Cell Surface Biotinylation Assay to Reveal Membrane-Associated Neuronal Cues: Negr1 Regulates Dendritic Arborization. Mol. Cell. Proteomics 2014, 13 (3), 733–748.

28. Nishimura, N.; Sasaki, T. Cell-Surface Biotinylation to Study Endocytosis and Recycling of Occludin. In Methods in Molecular Biology; 2008; pp 89–96.

29. Shin, B. K.; Wang, H.; Yim, A. M.; Le Naour, F.; Brichory, F.; Jang, J. H.; Zhao, R.; Puravs, E.; Tra, J.; Michael, C. W.; Misek, D. E.; Hanash, S. M. Global Profiling of the Cell Surface Proteome of Cancer Cells Uncovers an Abundance of Proteins with Chaperone Function. J. Biol. Chem. 2003, 278 (9), 7607–7616.

30. Wollscheid, B.; Bausch-Fluck, D.; Henderson, C.; O’Brien, R.; Bibel, M.; Schiess, R.; Aebersold, R.; Watts, J. D. Mass-Spectrometric Identification and Relative Quantification of N-Linked Cell Surface Glycoproteins. Nat. Biotechnol. 2009, 27 (4), 378–386.

31. Hofmann, A.; Bausch-Fluck, D.; Wollscheid, B. CSC Technology: Selective Labeling of Glycoproteins by Mild Oxidation to Phenotype Cells. Methods Mol. Biol. 2013, 951, 33–43.

32. Bausch-Fluck, D.; Hofmann, A.; Bock, T.; Frei, A. P.; Cerciello, F.; Jacobs, A.; Moest, H.; Omasits, U.; Gundry, R. L.; Yoon, C.; Schiess, R.; Schmidt, A.; Mirkowska, P.; Härtlová, A.; Van Eyk, J. E.; Bourquin, J. P.; Aebersold, R.; Boheler, K. R.; Zandstra, P.; Wollscheid, B. A Mass Spectrometric-Derived Cell Surface Protein Atlas. PLoS One 2015, 10 (4). 10.1371/journal.pone.0121314.

33. van Oostrum, M.; Müller, M.; Klein, F.; Bruderer, R.; Zhang, H.; Pedrioli, P. G. A.; Reiter, L.; Tsapogas, P.; Rolink, A.; Wollscheid, B. Classification of Mouse B Cell Types Using Surfaceome Proteotype Maps. Nat. Commun. 2019, 10 (1), 5734.

34. Gundry, R. L.; Riordon, D. R.; Tarasova, Y.; Chuppa, S.; Bhattacharya, S.; Juhasz, O.; Wiedemeier, O.; Milanovich, S.; Noto, F. K.; Tchernyshyov, I.; Raginski, K.; Bausch-Fluck, D.; Tae, H.-J.; Marshall, S.; Duncan, S. A.; Wollscheid, B.; Wersto, R. P.; Rao, S.; Van Eyk, J. E.; Boheler, K. R. A Cell Surfaceome Map for Immunophenotyping and Sorting Pluripotent Stem Cells. Mol. Cell. Proteomics 2012, 11 (8), 303–316.

35. Mallanna, S. K.; Waas, M.; Duncan, S. A.; Gundry, R. L. N-Glycoprotein Surfaceome of Human Induced Pluripotent Stem Cell Derived Hepatic Endoderm. Proteomics 2017, 17 (5). 10.1002/pmic.201600397.

36. Bausch-Fluck, D.; Hofmann, A.; Wollscheid, B. Cell Surface Capturing Technologies for the Surfaceome Discovery of Hepatocytes. In Methods in Molecular Biology; 2012; pp 1– 16.

37. Haverland, N. A.; Waas, M.; Ntai, I.; Keppel, T.; Gundry, R. L.; Kelleher, N. L. Cell Surface Proteomics of N-Linked Glycoproteins for Typing of Human Lymphocytes. Proteomics 2017, 17 (19), 1700156.

38. Weekes, M. P.; Antrobus, R.; Lill, J. R.; Duncan, L. M.; Hör, S.; Lehner, P. J. Comparative Analysis of Techniques to Purify Plasma Membrane Proteins. J. Biomol. Tech. 2010, 21 (3), 108–115.

39. Hörmann, K.; Stukalov, A.; Müller, A. C.; Heinz, L. X.; Superti-Furga, G.; Colinge, J.; Bennett, K. L. A Surface Biotinylation Strategy for Reproducible Plasma Membrane Protein Purification and Tracking of Genetic and Drug-Induced Alterations. J. Proteome Res. 2016, 15 (2), 647–658.

40. Gahmberg, C. G.; Tolvanen, M. Why Mammalian Cell Surface Proteins Are Glycoproteins. Trends Biochem. Sci. 1996, 21 (8), 308–311.

41. Hofmann, A.; Gerrits, B.; Schmidt, A.; Bock, T.; Bausch-Fluck, D.; Aebersold, R.; Wollscheid, B. Proteomic Cell Surface Phenotyping of Differentiating Acute Myeloid Leukemia Cells. Blood 2010, 116 (13), e26–e34.

42. Oldham, R. A. A.; Faber, M. L.; Keppel, T. R.; Buchberger, A. R.; Waas, M.; Hari, P.; Gundry, R. L.; Medin, J. A. Discovery and Validation of Surface N -Glycoproteins in MM Cell Lines and Patient Samples Uncovers Immunotherapy Targets. J. Immunother. Cancer 2020, 8 (2), e000915.

43. Boysen, G.; Bausch-Fluck, D.; Thoma, C. R.; Nowicka, A. M.; Stiehl, D. P.; Cima, I.; Luu, V.-D.; von Teichman, A.; Hermanns, T.; Sulser, T.; Ingold-Heppner, B.; Fankhauser, N.; Wenger, R. H.; Krek, W.; Krek, P.; Wollscheid, B.; Moch, H. Identification and Functional Characterization of PVHL-Dependent Cell Surface Proteins in Renal Cell Carcinoma. Neoplasia 2012, 14 (6), 535-IN17.

44. S Haun, R.; Yang Fan, C. CD109 Overexpression in Pancreatic Cancer Identified by Cell-Surface Glycoprotein Capture. J. Proteomics Bioinform. 2014, 01 (s10). 10.4172/jpb.S10-003.

45. Yan, M.; Yang, X.; Wang, L.; Clark, D.; Zuo, H.; Ye, D.; Chen, W.; Zhang, P. Plasma Membrane Proteomics of Tumor Spheres Identify CD166 as a Novel Marker for Cancer Stem-like Cells in Head and Neck Squamous Cell Carcinoma. Mol. Cell. Proteomics 2013, 12 (11), 3271–3284.

46. Kopecka, J.; Campia, I.; Jacobs, A.; Frei, A. P.; Ghigo, D.; Wollscheid, B.; Riganti, C. Carbonic Anhydrase XII Is a New Therapeutic Target to Overcome Chemoresistance in Cancer Cells. Oncotarget 2015, 6 (9), 6776–6793.

47. Go, Y.-M.; Chandler, J. D.; Jones, D. P. The Cysteine Proteome. Free Radic. Biol. Med. 2015, 84, 227–245.

48. Yamamoto, T.; Bishop, R. W.; Brown, M. S.; Goldstein, J. L.; Russell, D. W. Deletion in Cysteine-Rich Region of LDL Receptor Impedes Transport to Cell Surface in WHHL Rabbit. Science (80-.). 1986. 10.1126/science.3010466.

49. Iba, K.; Albrechtsen, R.; Gilpin, B. J.; Loechel, F.; Wewer, U. M. Cysteine-Rich Domain of Human ADAM 12 (Meltrin α) Supports Tumor Cell Adhesion. Am. J. Pathol. 1999, 154 (5), 1489–1501.

50. Ennion, S. J.; Evans, R. J. Conserved Cysteine Residues in the Extracellular Loop of the Human P2X 1 Receptor Form Disulfide Bonds and Are Involved in Receptor Trafficking to the Cell Surface. Mol. Pharmacol. 2002, 61 (2), 303–311.

51. Grzeszkiewicz, T. M.; Lindner, V.; Chen, N.; Lam, S. C. T.; Lau, L. F. The Angiogenic Factor Cysteine-Rich 61 (CYR61, CCN1) Supports Vascular Smooth Muscle Cell Adhesion and Stimulates Chemotaxis through Integrin Α6β1 and Cell Surface Heparan Sulfate Proteoglycans. Endocrinology 2002, 143 (4), 1441–1450.

52. Firsov, D.; Robert-Nicoud, M.; Gruender, S.; Schild, L.; Rossier, B. C. Mutational Analysis of Cysteine-Rich Domains of the Epithelium Sodium Channel (ENaC). J. Biol. Chem. 1999, 274 (5), 2743–2749.

53. Bestetti, S.; Medraño-Fernandez, I.; Galli, M.; Ghitti, M.; Bienert, G. P.; Musco, G.; Orsi, A.; Rubartelli, A.; Sitia, R. A Persulfidation-Based Mechanism Controls Aquaporin-8 Conductance. Sci. Adv. 2018, 4 (5). 10.1126/sciadv.aar5770.

54. Fiuza, C.; Bustin, M.; Talwar, S.; Tropea, M.; Gerstenberger, E.; Shelhamer, J. H.; Suffredini, A. F. Inflammation-Promoting Activity of HMGB1 on Human Microvascular Endothelial Cells. Blood 2003, 101 (7), 2652–2660.

55. Venereau, E.; Casalgrandi, M.; Schiraldi, M.; Antoine, D. J.; Cattaneo, A.; De Marchis, F.; Liu, J.; Antonelli, A.; Preti, A.; Raeli, L.; Shams, S. S.; Yang, H.; Varani, L.; Andersson, U.; Tracey, K. J.; Bachi, A.; Uguccioni, M.; Bianchi, M. E. Mutually Exclusive Redox Forms of HMGB1 Promote Cell Recruitment or Proinflammatory Cytokine Release. J. Exp. Med. 2012, 209 (9), 1519–1528.

56. Shi, C.; Zhang, Q.; Yao, Y.; Zeng, F.; Du, C.; Nijiati, S.; Wen, X.; Zhang, X.; Yang, H.; Chen, H.; Guo, Z.; Zhang, X.; Gao, J.; Guo, W.; Chen, X.; Zhou, Z. Targeting the Activity of T Cells by Membrane Surface Redox Regulation for Cancer Theranostics. Nat. Nanotechnol. 2023, 18 (1), 86–97.

57. Pellom, S. T.; Michalek, R. D.; Crump, K. E.; Langston, P. K.; Juneau, D. G.; Grayson, J. M. Increased Cell Surface Free Thiols Identify Effector CD8+ T Cells Undergoing T Cell Receptor Stimulation. PLoS One 2013, 8 (11), e81134.

58. Banerjee, R. Redox Outside the Box: Linking Extracellular Redox Remodeling with Intracellular Redox Metabolism. J. Biol. Chem. 2012, 287 (7), 4397–4402.

59. Ryser, H. J. P.; Levy, E. M.; Mandel, R.; DiSciullo, G. J. Inhibition of Human Immunodeficiency Virus Infection by Agents That Interfere with Thiol-Disulfide Interchange upon Virus-Receptor Interaction. Proc. Natl. Acad. Sci. 1994, 91 (10), 4559–4563.

60. Gallina, A.; Hanley, T. M.; Mandel, R.; Trahey, M.; Broder, C. C.; Viglianti, G. A.; Ryser, H. J. P. Inhibitors of Protein-Disulfide Isomerase Prevent Cleavage of Disulfide Bonds in Receptor-Bound Glycoprotein 120 and Prevent HIV-1 Entry. J. Biol. Chem. 2002, 277 (52), 50579–50588.

61. Li, D.; Ambrogio, L.; Shimamura, T.; Kubo, S.; Takahashi, M.; Chirieac, L. R.; Padera, R. F.; Shapiro, G. I.; Baum, A.; Himmelsbach, F.; Rettig, W. J.; Meyerson, M.; Solca, F.; Greulich, H.; Wong, K.-K. BIBW2992, an Irreversible EGFR/HER2 Inhibitor Highly Effective in Preclinical Lung Cancer Models. Oncogene 2008, 27 (34), 4702–4711.

62. Pan, Z.; Scheerens, H.; Li, S.-J.; Schultz, B. E.; Sprengeler, P. A.; Burrill, L. C.; Mendonca, R. V.; Sweeney, M. D.; Scott, K. C. K.; Grothaus, P. G.; Jeffery, D. A.; Spoerke, J. M.; Honigberg, L. A.; Young, P. R.; Dalrymple, S. A.; Palmer, J. T. Discovery of Selective Irreversible Inhibitors for Bruton’s Tyrosine Kinase. ChemMedChem 2007, 2 (1), 58–61.

63. Boatner, L. M.; Palafox, M. F.; Schweppe, D. K.; Backus, K. M. CysDB: A Human Cysteine Database Based on Experimental Quantitative Chemoproteomics. Cell Chem. Biol. 2023, 30 (6), 683–698.e3.

64. Yan, T.; Desai, H. S.; Boatner, L. M.; Yen, S. L.; Cao, J.; Palafox, M. F.; Jami-Alahmadi, Y.; Backus, K. M. SP3-FAIMS Chemoproteomics for High-Coverage Profiling of the Human Cysteinome**. ChemBioChem 2021, 22 (10), 1841–1851.

65. Desai, H. S.; Yan, T.; Backus, K. M. SP3-FAIMS-Enabled High-Throughput Quantitative Profiling of the Cysteinome. Curr. Protoc. 2022, 2 (7). 10.1002/cpz1.492.

66. Backus, K. M.; Correia, B. E.; Lum, K. M.; Forli, S.; Horning, B. D.; González-Páez, G. E.; Chatterjee, S.; Lanning, B. R.; Teijaro, J. R.; Olson, A. J.; Wolan, D. W.; Cravatt, B. F. Proteome-Wide Covalent Ligand Discovery in Native Biological Systems. Nature 2016, 534 (7608), 570–574.

67. Yan, T.; Palmer, A. B.; Geiszler, D. J.; Polasky, D. A.; Boatner, L. M.; Burton, N. R.; Armenta, E.; Nesvizhskii, A. I.; Backus, K. M. Enhancing Cysteine Chemoproteomic Coverage through Systematic Assessment of Click Chemistry Product Fragmentation. Anal. Chem. 2022, 94 (9), 3800–3810.

68. Jaffrey, S. R.; Snyder, S. H. The Biotin Switch Method for the Detection of S -Nitrosylated Proteins. Sci. STKE 2001, 2001 (86). 10.1126/stke.2001.86.pl1.

69. Leichert, L. I.; Gehrke, F.; Gudiseva, H. V.; Blackwell, T.; Ilbert, M.; Walker, A. K.; Strahler, J. R.; Andrews, P. C.; Jakob, U. Quantifying Changes in the Thiol Redox Proteome upon Oxidative Stress in Vivo. Proc. Natl. Acad. Sci. 2008, 105 (24), 8197–8202.

70. Desai, H. S.; Yan, T.; Yu, F.; Sun, A. W.; Villanueva, M.; Nesvizhskii, A. I.; Backus, K. M. SP3-Enabled Rapid and High Coverage Chemoproteomic Identification of Cell-State–Dependent Redox-Sensitive Cysteines. Mol. Cell. Proteomics 2022, 21 (4), 100218.

71. Fu, L.; Li, Z.; Liu, K.; Tian, C.; He, J.; He, J.; He, F.; Xu, P.; Yang, J. A Quantitative Thiol Reactivity Profiling Platform to Analyze Redox and Electrophile Reactive Cysteine Proteomes. Nat. Protoc. 2020, 15 (9), 2891–2919.

72. Xiao, H.; Jedrychowski, M. P.; Schweppe, D. K.; Huttlin, E. L.; Yu, Q.; Heppner, D. E.; Li, J.; Long, J.; Mills, E. L.; Szpyt, J.; He, Z.; Du, G.; Garrity, R.; Reddy, A.; Vaites, L. P.; Paulo, J. A.; Zhang, T.; Gray, N. S.; Gygi, S. P.; Chouchani, E. T. A Quantitative Tissue-Specific Landscape of Protein Redox Regulation during Aging. Cell 2020, 180 (5), 968–983.e24.

73. Yan, T.; Julio, A. R.; Villanueva, M.; Jones, A. E.; Ball, A. B.; Boatner, L. M.; Turmon, A. C.; Nguyễn, K. B.; Yen, S. L.; Desai, H. S.; Divakaruni, A. S.; Backus, K. M. Proximity-Labeling Chemoproteomics Defines the Subcellular Cysteinome and Inflammation-Responsive Mitochondrial Redoxome. Cell Chem. Biol. 2023, 30 (7), 811–827.e7.

74. Li, J.; Han, S.; Li, H.; Udeshi, N. D.; Svinkina, T.; Mani, D. R.; Xu, C.; Guajardo, R.; Xie, Q.; Li, T.; Luginbuhl, D. J.; Wu, B.; McLaughlin, C. N.; Xie, A.; Kaewsapsak, P.; Quake, S. R.; Carr, S. A.; Ting, A. Y.; Luo, L. Cell-Surface Proteomic Profiling in the Fly Brain Uncovers Wiring Regulators. Cell 2020, 180 (2), 373–386.e15.

75. Hughes, C. S.; Moggridge, S.; Müller, T.; Sorensen, P. H.; Morin, G. B.; Krijgsveld, J. Single-Pot, Solid-Phase-Enhanced Sample Preparation for Proteomics Experiments. Nat. Protoc. 2019, 14 (1), 68–85.

76. Cao, J.; Boatner, L. M.; Desai, H. S.; Burton, N. R.; Armenta, E.; Chan, N. J.; Castellón, J. O.; Backus, K. M. Multiplexed CuAAC Suzuki–Miyaura Labeling for Tandem Activity-Based Chemoproteomic Profiling. Anal. Chem. 2021, 93 (4), 2610–2618.

77. Sielaff, M.; Kuharev, J.; Bohn, T.; Hahlbrock, J.; Bopp, T.; Tenzer, S.; Distler, U. Evaluation of FASP, SP3, and IST Protocols for Proteomic Sample Preparation in the Low Microgram Range. J. Proteome Res. 2017, 16 (11), 4060–4072.

78. Thul, P. J.; Åkesson, L.; Wiking, M.; Mahdessian, D.; Geladaki, A.; Ait Blal, H.; Alm, T.; Asplund, A.; Björk, L.; Breckels, L. M.; Bäckström, A.; Danielsson, F.; Fagerberg, L.; Fall, J.; Gatto, L.; Gnann, C.; Hober, S.; Hjelmare, M.; Johansson, F.; Lee, S.; Lindskog, C.; Mulder, J.; Mulvey, C. M.; Nilsson, P.; Oksvold, P.; Rockberg, J.; Schutten, R.; Schwenk, J. M.; Sivertsson, Å.; Sjöstedt, E.; Skogs, M.; Stadler, C.; Sullivan, D. P.; Tegel, H.; Winsnes, C.; Zhang, C.; Zwahlen, M.; Mardinoglu, A.; Pontén, F.; von Feilitzen, K.; Lilley, K. S.; Uhlén, M.; Lundberg, E. A Subcellular Map of the Human Proteome. Science (80-.). 2017, 356 (6340). 10.1126/science.aal3321.

79. Bateman, A.; Martin, M.-J.; Orchard, S.; Magrane, M.; Ahmad, S.; Alpi, E.; Bowler-Barnett, E. H.; Britto, R.; Bye-A-Jee, H.; Cukura, A.; Denny, P.; Dogan, T.; Ebenezer, T.; Fan, J.; Garmiri, P.; da Costa Gonzales, L. J.; Hatton-Ellis, E.; Hussein, A.; Ignatchenko, A.; Insana, G.; Ishtiaq, R.; Joshi, V.; Jyothi, D.; Kandasaamy, S.; Lock, A.; Luciani, A.; Lugaric, M.; Luo, J.; Lussi, Y.; MacDougall, A.; Madeira, F.; Mahmoudy, M.; Mishra, A.; Moulang, K.; Nightingale, A.; Pundir, S.; Qi, G.; Raj, S.; Raposo, P.; Rice, D. L.; Saidi, R.; Santos, R.; Speretta, E.; Stephenson, J.; Totoo, P.; Turner, E.; Tyagi, N.; Vasudev, P.; Warner, K.; Watkins, X.; Zaru, R.; Zellner, H.; Bridge, A. J.; Aimo, L.; Argoud-Puy, G.; Auchincloss, A. H.; Axelsen, K. B.; Bansal, P.; Baratin, D.; Batista Neto, T. M.; Blatter, M.-C.; Bolleman, J. T.; Boutet, E.; Breuza, L.; Gil, B. C.; Casals-Casas, C.; Echioukh, K. C.; Coudert, E.; Cuche, B.; de Castro, E.; Estreicher, A.; Famiglietti, M. L.; Feuermann, M.; Gasteiger, E.; Gaudet, P.; Gehant, S.; Gerritsen, V.; Gos, A.; Gruaz, N.; Hulo, C.; Hyka-Nouspikel, N.; Jungo, F.; Kerhornou, A.; Le Mercier, P.; Lieberherr, D.; Masson, P.; Morgat, A.; Muthukrishnan, V.; Paesano, S.; Pedruzzi, I.; Pilbout, S.; Pourcel, L.; Poux, S.; Pozzato, M.; Pruess, M.; Redaschi, N.; Rivoire, C.; Sigrist, C. J. A.; Sonesson, K.; Sundaram, S.; Wu, C. H.; Arighi, C. N.; Arminski, L.; Chen, C.; Chen, Y.; Huang, H.; Laiho, K.; McGarvey, P.; Natale, D. A.; Ross, K.; Vinayaka, C. R.; Wang, Q.; Wang, Y.; Zhang, J. UniProt: The Universal Protein Knowledgebase in 2023. Nucleic Acids Res. 2023, 51 (D1), D523–D531.

80. Zhu, L.; Malatras, A.; Thorley, M.; Aghoghogbe, I.; Mer, A.; Duguez, S.; Butler-Browne, G.; Voit, T.; Duddy, W. CellWhere: Graphical Display of Interaction Networks Organized on Subcellular Localizations. Nucleic Acids Res. 2015, 43 (W1), W571–W575.

81. Yan, T.; Julio, A. R.; Villanueva, M.; Jones, A. E.; Ball, A. A. B.; Boatner, L. M.; Turmon, A. C.; Yen, S. L.; Desai, H. S.; Divakaruni, A. S.; Backus, K. M. Proximity-Labeling Chemoproteomics Defines the Subcellular Cysteinome and Inflammation-Responsive Mitochondrial Redoxome. bioRxiv Prepr. Serv. Biol. 2023. 10.1101/2023.01.22.525042.

82. Weerapana, E.; Wang, C.; Simon, G. M.; Richter, F.; Khare, S.; Dillon, M. B. D.; Bachovchin, D. A.; Mowen, K.; Baker, D.; Cravatt, B. F. Quantitative Reactivity Profiling Predicts Functional Cysteines in Proteomes. Nature 2010, 468 (7325), 790–797.

83. Yu, F.; Haynes, S. E.; Nesvizhskii, A. I. IonQuant Enables Accurate and Sensitive Label-Free Quantification With FDR-Controlled Match-Between-Runs. Mol. Cell. Proteomics 2021, 20, 100077.

84. Guo, H.; Wang, J.; Ren, S.; Zheng, L.-F.; Zhuang, Y.-X.; Li, D.-L.; Sun, H.-H.; Liu, L.-Y.; Xie, C.; Wu, Y.-Y.; Wang, H.-R.; Deng, X.; Li, P.; Zhao, T.-J. Targeting EGFR-Dependent Tumors by Disrupting an ARF6-Mediated Sorting System. Nat. Commun. 2022, 13 (1), 6004.

85. Runkle, K. B.; Kharbanda, A.; Stypulkowski, E.; Cao, X.-J.; Wang, W.; Garcia, B. A.; Witze, E. S. Inhibition of DHHC20-Mediated EGFR Palmitoylation Creates a Dependence on EGFR Signaling. Mol. Cell 2016, 62 (3), 385–396.

86. Kuleshov, M. V.; Jones, M. R.; Rouillard, A. D.; Fernandez, N. F.; Duan, Q.; Wang, Z.; Koplev, S.; Jenkins, S. L.; Jagodnik, K. M.; Lachmann, A.; McDermott, M. G.; Monteiro, C. D.; Gundersen, G. W.; Ma’ayan, A. Enrichr: A Comprehensive Gene Set Enrichment Analysis Web Server 2016 Update. Nucleic Acids Res. 2016, 44 (W1), W90–W97.

87. Paiva, B.; Gutiérrez, N.-C.; Chen, X.; Vídriales, M.-B.; Montalbán, M.-Á.; Rosiñol, L.; Oriol, A.; Martínez-López, J.; Mateos, M.-V.; López-Corral, L.; Díaz-Rodríguez, E.; Pérez, J.-J.; Fernández-Redondo, E.; de Arriba, F.; Palomera, L.; Bengoechea, E.; Terol, M.-J.; de Paz, R.; Martin, A.; Hernández, J.; Orfao, A.; Lahuerta, J.-J.; Bladé, J.; Pandiella, A.; Miguel, J.-F. S. Clinical Significance of CD81 Expression by Clonal Plasma Cells in High-Risk Smoldering and Symptomatic Multiple Myeloma Patients. Leukemia 2012, 26 (8), 1862–1869.

88. Alam, A.; Woo, J.-S.; Schmitz, J.; Prinz, B.; Root, K.; Chen, F.; Bloch, J. S.; Zenobi, R.; Locher, K. P. Structural Basis of Transcobalamin Recognition by Human CD320 Receptor. Nat. Commun. 2016, 7 (1), 12100.

89. Jumper, J.; Evans, R.; Pritzel, A.; Green, T.; Figurnov, M.; Ronneberger, O.; Tunyasuvunakool, K.; Bates, R.; Žídek, A.; Potapenko, A.; Bridgland, A.; Meyer, C.; Kohl, S. A. A.; Ballard, A. J.; Cowie, A.; Romera-Paredes, B.; Nikolov, S.; Jain, R.; Adler, J.; Back, T.; Petersen, S.; Reiman, D.; Clancy, E.; Zielinski, M.; Steinegger, M.; Pacholska, M.; Berghammer, T.; Bodenstein, S.; Silver, D.; Vinyals, O.; Senior, A. W.; Kavukcuoglu, K.; Kohli, P.; Hassabis, D. Highly Accurate Protein Structure Prediction with AlphaFold. Nature 2021, 596 (7873), 583–589.

90. Varadi, M.; Anyango, S.; Deshpande, M.; Nair, S.; Natassia, C.; Yordanova, G.; Yuan, D.; Stroe, O.; Wood, G.; Laydon, A.; Žídek, A.; Green, T.; Tunyasuvunakool, K.; Petersen, S.; Jumper, J.; Clancy, E.; Green, R.; Vora, A.; Lutfi, M.; Figurnov, M.; Cowie, A.; Hobbs, N.; Kohli, P.; Kleywegt, G.; Birney, E.; Hassabis, D.; Velankar, S. AlphaFold Protein Structure Database: Massively Expanding the Structural Coverage of Protein-Sequence Space with High-Accuracy Models. Nucleic Acids Res. 2022, 50 (D1), D439–D444.

91. Zhang, Z.; Rohweder, P. J.; Ongpipattanakul, C.; Basu, K.; Bohn, M.-F.; Dugan, E. J.; Steri, V.; Hann, B.; Shokat, K. M.; Craik, C. S. A Covalent Inhibitor of K-Ras(G12C) Induces MHC Class I Presentation of Haptenated Peptide Neoepitopes Targetable by Immunotherapy. Cancer Cell 2022, 40 (9), 1060–1069.e7.

92. Hattori, T.; Maso, L.; Araki, K. Y.; Koide, A.; Hayman, J.; Akkapeddi, P.; Bang, I.; Neel, B. G.; Koide, S. Creating MHC-Restricted Neoantigens with Covalent Inhibitors That Can Be Targeted by Immune Therapy. Cancer Discov. 2023, 13 (1), 132–145.

93. Cravatt, B. F.; Wright, A. T.; Kozarich, J. W. Activity-Based Protein Profiling: From Enzyme Chemistry to Proteomic Chemistry. Annu. Rev. Biochem. 2008, 77 (1), 383–414.

94. Zamorano Cuervo, N.; Fortin, A.; Caron, E.; Chartier, S.; Grandvaux, N. Pinpointing Cysteine Oxidation Sites by High-Resolution Proteomics Reveals a Mechanism of Redox-Dependent Inhibition of Human STING. Sci. Signal. 2021, 14 (680). 10.1126/scisignal.aaw4673.

95. Sachs, A.; Moore, E.; Kosaloglu-Yalcin, Z.; Peters, B.; Sidney, J.; Rosenberg, S. A.; Robbins, P. F.; Sette, A. Impact of Cysteine Residues on MHC Binding Predictions and Recognition by Tumor-Reactive T Cells. J. Immunol. 2020, 205 (2), 539–549.

96. Abo, M.; Weerapana, E. Chemical Probes for Redox Signaling and Oxidative Stress. Antioxid. Redox Signal. 2019, 30 (10), 1369–1386.

97. Sethuraman, M.; McComb, M. E.; Heibeck, T.; Costello, C. E.; Cohen, R. A. Isotope-Coded Affinity Tag Approach to Identify and Quantify Oxidant-Sensitive Protein Thiols. Mol. Cell. Proteomics 2004, 3 (3), 273–278.

98. Gygi, S. P.; Rist, B.; Gerber, S. A.; Turecek, F.; Gelb, M. H.; Aebersold, R. Quantitative Analysis of Complex Protein Mixtures Using Isotope-Coded Affinity Tags. Nat. Biotechnol. 1999, 17 (10), 994–999.

99. Abo, M.; Li, C.; Weerapana, E. Isotopically-Labeled Iodoacetamide-Alkyne Probes for Quantitative Cysteine-Reactivity Profiling. Mol. Pharm. 2018, 15 (3), 743–749.

100. Shakir, S.; Vinh, J.; Chiappetta, G. Quantitative Analysis of the Cysteine Redoxome by Iodoacetyl Tandem Mass Tags. Anal. Bioanal. Chem. 2017, 409 (15), 3821–3830.

101. Vajrychova, M.; Salovska, B.; Pimkova, K.; Fabrik, I.; Tambor, V.; Kondelova, A.; Bartek, J.; Hodny, Z. Quantification of Cellular Protein and Redox Imbalance Using SILAC-IodoTMT Methodology. Redox Biol. 2019. 10.1016/j.redox.2019.101227.

102. Franchina, D. G.; Dostert, C.; Brenner, D. Reactive Oxygen Species: Involvement in T Cell Signaling and Metabolism. Trends Immunol. 2018, 39 (6), 489–502.

103. Cemerski, S.; Cantagrel, A.; van Meerwijk, J. P. M.; Romagnoli, P. Reactive Oxygen Species Differentially Affect T Cell Receptor-Signaling Pathways*. J. Biol. Chem. 2002, 277 (22), 19585–19593.

104. Belikov, A. V.; Schraven, B.; Simeoni, L. T Cells and Reactive Oxygen Species. J. Biomed. Sci. 2015, 22 (1), 85.

105. Vinogradova, E. V.; Zhang, X.; Remillard, D.; Lazar, D. C.; Suciu, R. M.; Wang, Y.; Bianco, G.; Yamashita, Y.; Crowley, V. M.; Schafroth, M. A.; Yokoyama, M.; Konrad, D. B.; Lum, K. M.; Simon, G. M.; Kemper, E. K.; Lazear, M. R.; Yin, S.; Blewett, M. M.; Dix, M. M.; Nguyen, N.; Shokhirev, M. N.; Chin, E. N.; Lairson, L. L.; Melillo, B.; Schreiber, S. L.; Forli, S.; Teijaro, J. R.; Cravatt, B. F. An Activity-Guided Map of Electrophile-Cysteine Interactions in Primary Human T Cells. Cell 2020, 182 (4), 1009–1026.e29.

106. Trujillo, J. A.; Croft, N. P.; Dudek, N. L.; Channappanavar, R.; Theodossis, A.; Webb, A. I.; Dunstone, M. A.; Illing, P. T.; Butler, N. S.; Fett, C.; Tscharke, D. C.; Rossjohn, J.; Perlman, S.; Purcell, A. W. The Cellular Redox Environment Alters Antigen Presentation. J. Biol. Chem. 2014, 289 (40), 27979–27991.

107. Yan, Z.; Garg, S. K.; Kipnis, J.; Banerjee, R. Extracellular Redox Modulation by Regulatory T Cells. Nat. Chem. Biol. 2009, 5 (10), 721–723.

108. Kesarwani, P.; Murali, A. K.; Al-Khami, A. A.; Mehrotra, S. Redox Regulation of T-Cell Function: From Molecular Mechanisms to Significance in Human Health and Disease. Antioxid. Redox Signal. 2013, 18 (12), 1497–1534.

109. Metcalfe, C.; Cresswell, P.; Ciaccia, L.; Thomas, B.; Barclay, A. N. Labile Disulfide Bonds Are Common at the Leucocyte Cell Surface. Open Biol. 2011, 1 (3), 110010.

110. Metcalfe, C.; Cresswell, P.; Barclay, A. N. Interleukin-2 Signalling Is Modulated by a Labile Disulfide Bond in the CD132 Chain of Its Receptor. Open Biol. 2012, 2 (1), 110036.

111. Yamamoto, T.; Bishop, R. W.; Brown, M. S.; Goldstein, J. L.; Russell, D. W. Deletion in Cysteine-Rich Region of LDL Receptor Impedes Transport to Cell Surface in WHHL Rabbit. Science (80-.). 1986, 232 (4755), 1230–1237.

112. Oliveira, M. I.; Gonçalves, C. M.; Pinto, M.; Fabre, S.; Santos, A. M.; Lee, S. F.; Castro, M. A. A.; Nunes, R. J.; Barbosa, R. R.; Parnes, J. R.; Yu, C.; Davis, S. J.; Moreira, A.; Bismuth, G.; Carmo, A. M. CD6 Attenuates Early and Late Signaling Events, Setting Thresholds for T-Cell Activation. Eur. J. Immunol. 2012, 42 (1), 195–205.

113. Chappell, P. E.; Garner, L. I.; Yan, J.; Metcalfe, C.; Hatherley, D.; Johnson, S.; Robinson, C. V.; Lea, S. M.; Brown, M. H. Structures of CD6 and Its Ligand CD166 Give Insight into Their Interaction. Structure 2015, 23 (8), 1426–1436.

114. Aertgeerts, K.; Ye, S.; Shi, L.; Prasad, S. G.; Witmer, D.; Chi, E.; Sang, B.-C.; Wijnands, R. A.; Webb, D. R.; Swanson, R. V. N-Linked Glycosylation of Dipeptidyl Peptidase IV (CD26): Effects on Enzyme Activity, Homodimer Formation, and Adenosine Deaminase Binding. Protein Sci. 2004, 13 (1), 145–154.

115. Ginés, S.; Mariño, M.; Mallol, J.; Canela, E. I.; Morimoto, C.; Callebaut, C.; Hovanessian, A.; Casadó, V.; Lluis, C.; Franco, R. Regulation of Epithelial and Lymphocyte Cell Adhesion by Adenosine Deaminase-CD26 Interaction. Biochem. J. 2002, 361 (Pt 2), 203–209.

116. Ikushima, H.; Munakata, Y.; Ishii, T.; Iwata, S.; Terashima, M.; Tanaka, H.; Schlossman, S. F.; Morimoto, C. Internalization of CD26 by Mannose 6-Phosphate/Insulin-like Growth Factor II Receptor Contributes to T Cell Activation. Proc. Natl. Acad. Sci. U. S. A. 2000, 97 (15), 8439–8444.

117. Durinx, C.; Lambeir, A. M.; Bosmans, E.; Falmagne, J. B.; Berghmans, R.; Haemers, A.; Scharpé, S.; De Meester, I. Molecular Characterization of Dipeptidyl Peptidase Activity in Serum: Soluble CD26/Dipeptidyl Peptidase IV Is Responsible for the Release of X-Pro Dipeptides. Eur. J. Biochem. 2000, 267 (17), 5608–5613.

118. Wu, L.; Fu, J.; Shen, S.-H. SKAP55 Coupled with CD45 Positively Regulates T-Cell Receptor-Mediated Gene Transcription. Mol. Cell. Biol. 2002, 22 (8), 2673–2686.

119. Nam, H.-J.; Poy, F.; Saito, H.; Frederick, C. A. Structural Basis for the Function and Regulation of the Receptor Protein Tyrosine Phosphatase CD45. J. Exp. Med. 2005, 201 (3), 441–452.

120. Oravecz, T.; Monostori, É.; Adrian, O.; István Andó, É. K. Novel Heterogeneity of the Leucocyte Common Antigen (CD45): Disulfide-Bound Heterodimers between CD45 and an 80 KDa Polypeptide. Immunol. Lett. 1994, 40 (1), 7–11.

121. Matthias, L. J.; Azimi, I.; Tabrett, C. A.; Hogg, P. J. Reduced Monomeric CD4 Is the Preferred Receptor for HIV. J. Biol. Chem. 2010, 285 (52), 40793–40799.

122. Cerutti, N.; Killick, M.; Jugnarain, V.; Papathanasopoulos, M.; Capovilla, A. Disulfide Reduction in CD4 Domain 1 or 2 Is Essential for Interaction with HIV Glycoprotein 120 (Gp120), Which Impairs Thioredoxin-Driven CD4 Dimerization. J. Biol. Chem. 2014, 289 (15), 10455–10465.

123. Dong, G.; Wearsch, P. A.; Peaper, D. R.; Cresswell, P.; Reinisch, K. M. Insights into MHC Class I Peptide Loading from the Structure of the Tapasin-ERp57 Thiol Oxidoreductase Heterodimer. Immunity 2009, 30 (1), 21–32.

124. Katz, B. A.; Kossiakoff, A. The Crystallographically Determined Structures of Atypical Strained Disulfides Engineered into Subtilisin. J. Biol. Chem. 1986, 261 (33), 15480–15485.

125. Wells, J. A.; Powers, D. B. In Vivo Formation and Stability of Engineered Disulfide Bonds in Subtilisin. J. Biol. Chem. 1986, 261 (14), 6564–6570.

126. Wouters, M. A.; Lau, K. K.; Hogg, P. J. Cross-Strand Disulphides in Cell Entry Proteins: Poised to Act. BioEssays 2004, 26 (1), 73–79.

127. Wong, J. W. H.; Hogg, P. J. Allosteric Disulfide Bonds. In Folding of Disulfide Proteins; Springer New York: New York, NY, 2011; pp 151–182.

128. Ezeriņa, D.; Takano, Y.; Hanaoka, K.; Urano, Y.; Dick, T. P. N-Acetyl Cysteine Functions as a Fast-Acting Antioxidant by Triggering Intracellular H2S and Sulfane Sulfur Production. Cell Chem. Biol. 2018, 25 (4), 447–459.e4.

129. Mokhtari, V.; Afsharian, P.; Shahhoseini, M.; Kalantar, S. M.; Moini, A. A Review on Various Uses of N-Acetyl Cysteine. Cell J. 2017, 19 (1), 11–17.

130. Sahaf, B.; Heydari, K.; Herzenberg, L. A.; Herzenberg, L. A. Lymphocyte Surface Thiol Levels. Proc. Natl. Acad. Sci. 2003, 100 (7), 4001–4005.

131. Imamoto, Y.; Kataoka, M.; Tokunaga, F.; Palczewski, K. Light-Induced Conformational Changes of Rhodopsin Probed by Fluorescent Alexa594 Immobilized on the Cytoplasmic Surface. Biochemistry 2000, 39 (49), 15225–15233.

132. Hu, M.; Zhang, H.; Liu, Q.; Hao, Q. Structural Basis for Human PECAM-1-Mediated Trans-Homophilic Cell Adhesion. Sci. Rep. 2016, 6 (1), 38655.

133. Sachs, U. J. H.; Andrei-Selmer, C. L.; Maniar, A.; Weiss, T.; Paddock, C.; Orlova, V. V.; Choi, E. Y.; Newman, P. J.; Preissner, K. T.; Chavakis, T.; Santoso, S. The Neutrophil-Specific Antigen CD177 Is a Counter-Receptor for Platelet Endothelial Cell Adhesion Molecule-1 (CD31). J. Biol. Chem. 2007, 282 (32), 23603–23612.

134. Ymer, S. I.; Greenall, S. A.; Cvrljevic, A.; Cao, D. X.; Donoghue, J. F.; Epa, V. C.; Scott, A. M.; Adams, T. E.; Johns, T. G. Glioma Specific Extracellular Missense Mutations in the First Cysteine Rich Region of Epidermal Growth Factor Receptor (EGFR) Initiate Ligand Independent Activation. Cancers (Basel). 2011, 3 (2), 2032–2049.

135. Colby, S.; Yehia, L.; Niazi, F.; Chen, J.; Ni, Y.; Mester, J. L.; Eng, C. Exome Sequencing Reveals Germline Gain-of-Function EGFR Mutation in an Adult with Lhermitte–Duclos Disease. Mol. Case Stud. 2016, 2 (6), a001230.

136. Hobbs, H. H.; Brown, M. S.; Goldstein, J. L. Molecular Genetics of the LDL Receptor Gene in Familial Hypercholesterolemia. Hum. Mutat. 1992, 1 (6), 445–466.

137. Soutar, A. K.; Naoumova, R. P. Mechanisms of Disease: Genetic Causes of Familial Hypercholesterolemia. Nat. Clin. Pract. Cardiovasc. Med. 2007, 4 (4), 214–225.

138. Henrie, A.; Hemphill, S. E.; Ruiz-Schultz, N.; Cushman, B.; DiStefano, M. T.; Azzariti, D.; Harrison, S. M.; Rehm, H. L.; Eilbeck, K. ClinVar Miner: Demonstrating Utility of a Web-Based Tool for Viewing and Filtering ClinVar Data. Hum. Mutat. 2018, 39 (8), 1051–1060.

139. Benito-Vicente, A.; Uribe, K. B.; Siddiqi, H.; Jebari, S.; Galicia-Garcia, U.; Larrea-Sebal, A.; Cenarro, A.; Stef, M.; Ostolaza, H.; Civeira, F.; Palacios, L.; Martin, C. Replacement of Cysteine at Position 46 in the First Cysteine-Rich Repeat of the LDL Receptor Impairs Apolipoprotein Recognition. PLoS One 2018, 13 (10), e0204771.

140. Hansen, R. E.; Roth, D.; Winther, J. R. Quantifying the Global Cellular Thiol–Disulfide Status. Proc. Natl. Acad. Sci. 2009, 106 (2), 422–427.

141. Brandes, N.; Reichmann, D.; Tienson, H.; Leichert, L. I.; Jakob, U. Using Quantitative Redox Proteomics to Dissect the Yeast Redoxome. J. Biol. Chem. 2011, 286 (48), 41893– 41903.

142. Go, Y.-M.; Jones, D. P. Redox Compartmentalization in Eukaryotic Cells. Biochim. Biophys. Acta - Gen. Subj. 2008, 1780 (11), 1273–1290.

143. Jones, D. P.; Go, Y. -M. Redox Compartmentalization and Cellular Stress. Diabetes, Obes. Metab. 2010, 12 (s2), 116–125.

144. Li, H.; Remsberg, J. R.; Won, S. J.; Zhao, K. T.; Huang, T. P.; Lu, B.; Simon, G. M.; Liu, D. R.; Cravatt, B. F. Assigning Functionality to Cysteines by Base Editing of Cancer Dependency Genes. bioRxiv 2022.

145. Desai, H.; Ofori, S.; Boatner, L.; Yu, F.; Villanueva, M.; Ung, N.; Nesvizhskii, A. I.; Backus, K. Multi-Omic Stratification of the Missense Variant Cysteinome. bioRxiv Prepr. Serv. Biol. 2023. 10.1101/2023.08.12.553095.

146. Cole, K. S.; Grandjean, J. M. D.; Chen, K.; Witt, C. H.; O’Day, J.; Shoulders, M. D.; Wiseman, R. L.; Weerapana, E. Characterization of an A-Site Selective Protein Disulfide Isomerase A1 Inhibitor. Biochemistry 2018, 57 (13), 2035–2043.

147. Almasy, K. M.; Davies, J. P.; Lisy, S. M.; Tirgar, R.; Tran, S. C.; Plate, L. Small-Molecule Endoplasmic Reticulum Proteostasis Regulator Acts as a Broad-Spectrum Inhibitor of Dengue and Zika Virus Infections. Proc. Natl. Acad. Sci. 2021, 118 (3). 10.1073/pnas.2012209118.

148. Xie, Z.; Bailey, A.; Kuleshov, M. V.; Clarke, D. J. B.; Evangelista, J. E.; Jenkins, S. L.; Lachmann, A.; Wojciechowicz, M. L.; Kropiwnicki, E.; Jagodnik, K. M.; Jeon, M.; Ma’ayan, A. Gene Set Knowledge Discovery with Enrichr. Curr. Protoc. 2021, 1 (3). 10.1002/cpz1.90.

149. Deutsch, E. W.; Csordas, A.; Sun, Z.; Jarnuczak, A.; Perez-Riverol, Y.; Ternent, T.; Campbell, D. S.; Bernal-Llinares, M.; Okuda, S.; Kawano, S.; Moritz, R. L.; Carver, J. J.; Wang, M.; Ishihama, Y.; Bandeira, N.; Hermjakob, H.; Vizcaíno, J. A. The ProteomeXchange Consortium in 2017: Supporting the Cultural Change in Proteomics Public Data Deposition. Nucleic Acids Res. 2017, 45 (D1), D1100–D1106.

150. Perez-Riverol, Y.; Bai, J.; Bandla, C.; García-Seisdedos, D.; Hewapathirana, S.; Kamatchinathan, S.; Kundu, D. J.; Prakash, A.; Frericks-Zipper, A.; Eisenacher, M.; Walzer, M.; Wang, S.; Brazma, A.; Vizcaíno, J. A. The PRIDE Database Resources in 2022: A Hub for Mass Spectrometry-Based Proteomics Evidences. Nucleic Acids Res. 2022, 50 (D1), D543–D552.

151. Kong, A. T.; Leprevost, F. V.; Avtonomov, D. M.; Mellacheruvu, D.; Nesvizhskii, A. I. MSFragger: Ultrafast and Comprehensive Peptide Identification in Mass Spectrometry– Based Proteomics. Nat. Methods 2017, 14 (5), 513–520.

152. Shteynberg, D. D.; Deutsch, E. W.; Campbell, D. S.; Hoopmann, M. R.; Kusebauch, U.; Lee, D.; Mendoza, L.; Midha, M. K.; Sun, Z.; Whetton, A. D.; Moritz, R. L. PTMProphet: Fast and Accurate Mass Modification Localization for the Trans-Proteomic Pipeline. J. Proteome Res. 2019, 18 (12), 4262–4272.

