## Supplementary Information for "Defining the Cell Surface Cysteinome using Two-step Enrichment Proteomics"

[a]Department of Biological Chemistry, David Geffen School of Medicine, UCLA, Los Angeles, CA 90095 (USA); [b]Department of Chemistry and Biochemistry, UCLA, Los Angeles, CA 90095 (USA); [c]Department of Pathology and Laboratory Medicine, UCLA, Los Angeles, CA 90095, USA, [d]DOE Institute for Genomics and Proteomics, UCLA, Los Angeles, CA 90095 (USA); [e]Jonsson Comprehensive Cancer Center, UCLA, Los Angeles, CA 90095 (USA); [f]Eli and Edythe Broad Center of Regenerative Medicine and Stem Cell Research, UCLA, Los Angeles, CA 90095 (USA).

**Table of Contents**


| | | |
|---|---|---|
| **A.** | **Supplementary Figures** | **3–12** |
| **B.** | **Supplementary Tables** | **12–14** |

### A. Supplementary Figures

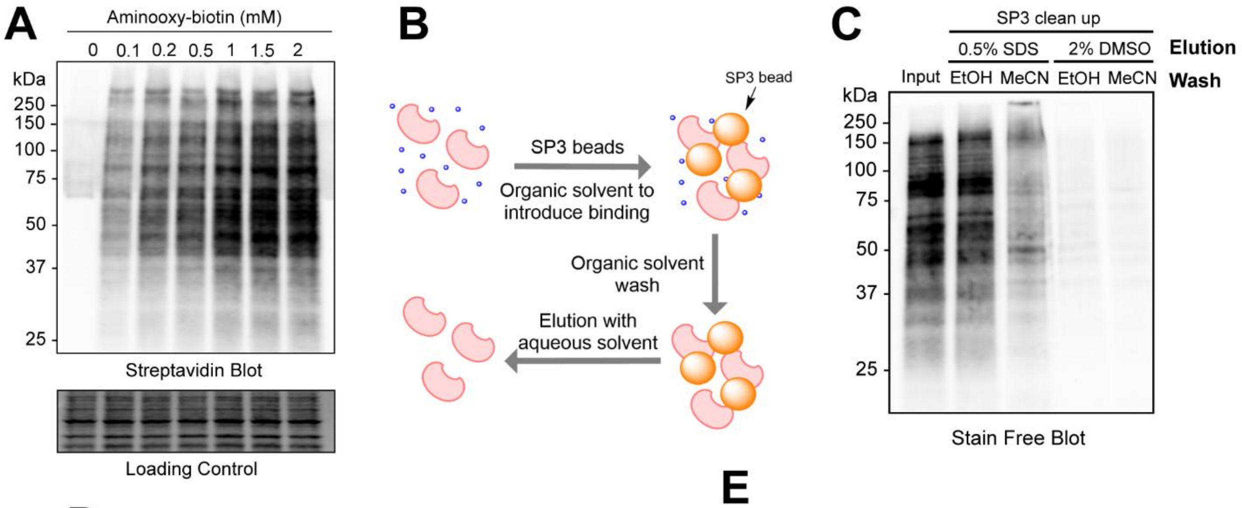


D

| Washing condition | Eluting condition | Protein recovery rate from 2mg/mL lysate |
|---|---|---|
| EtOH | 1% SDS/PBS | 65.5% |
| | 0.5% SDS/PBS | 70.3% |
| | 0.2% SDS/PBS | 78.7% |
| | 0.1% SDS/PBS | 68.0% |
| MeCN | 1% SDS/PBS | 8.4% |
| | 0.5% SDS/PBS | 6.2% |
| | 0.2% SDS/PBS | 8.1% |
| | 0.1% SDS/PBS | 8.4% |

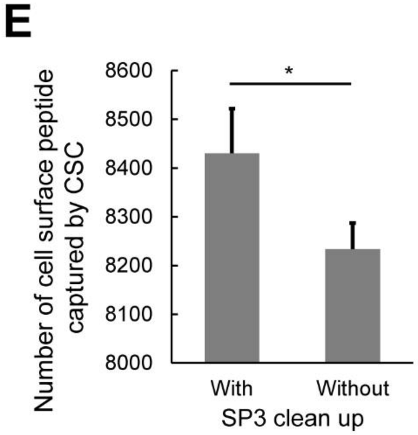


**Figure S1.** A) Streptavidin blot of Cell Surface Capture (CSC) with increasing amount of Aminooxy-biotin. B) Schematic workflow of protein level SP3 clean up. C) Stain free blot of the input and eluted lysates of SP3 clean up with different conditions. D) Protein recovery rate of SP3 clean up with different protein elution conditions. Protein concentration was determined by DC assay. E) Number of cell surface peptides was captured with or without SP3 clean up. Statistical significance was calculated with Student's t-tests, * $p$<0.05. Data in panel E are represented as mean ± stdev. MS experiments were conducted in 2 replicates.

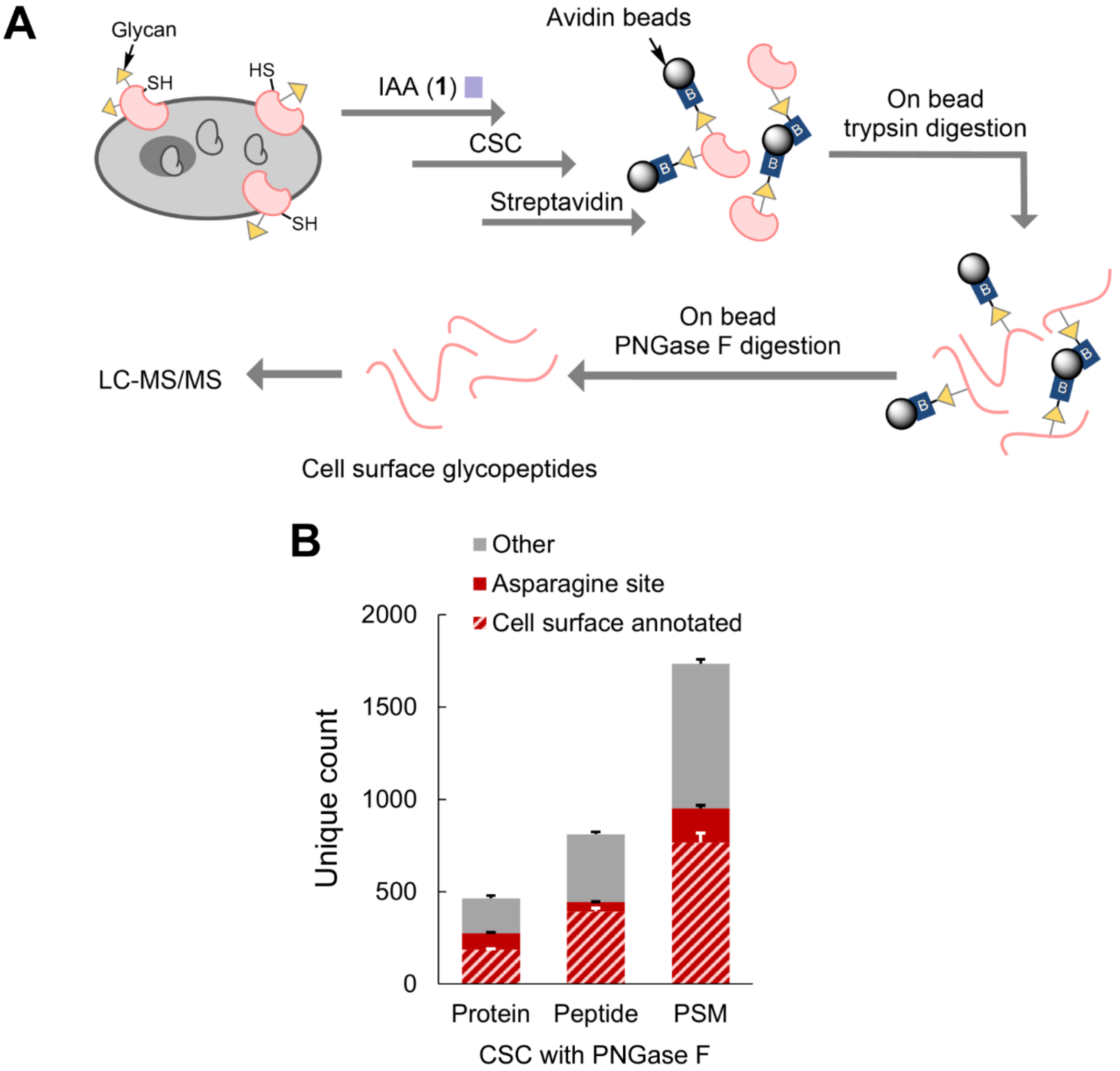


**Figure S2.** A) The schematic workflow of Cell Surface Capture (CSC) coupled with treatment of PNGase F. B) Unique counts of proteins, peptides and PSMs were identified with CSC with PNGase F. PNGase F treatment was for 24 h at 37 °C. Data in panel B are represented as mean ± stdev. MS experiments were conducted in 2 replicates in Jurkats cells.

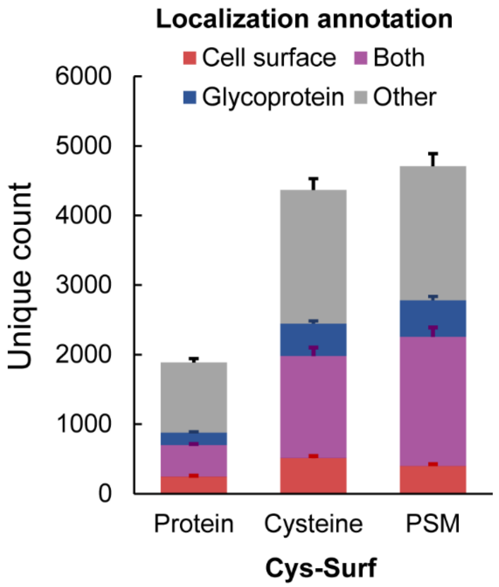


**Figure S3** Unique counts of proteins, cysteines and PSMs identified with Cys-Surf. Data in are represented as mean ± stdev. MS experiments were conducted in 3 replicates in Jurkats cells.

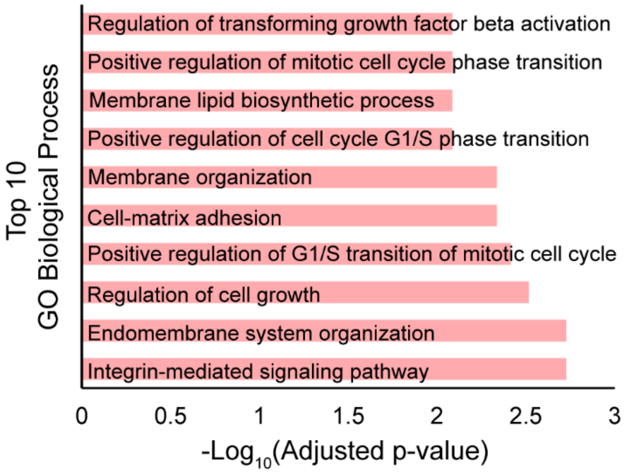


**Figure S4.** Top 10 GO biological processes enriched with proteins harboring ligandable cell surface cysteines ($Log_2(H/L) > 1$). MS experiments were conducted in 3 replicates in Jurkats cells.

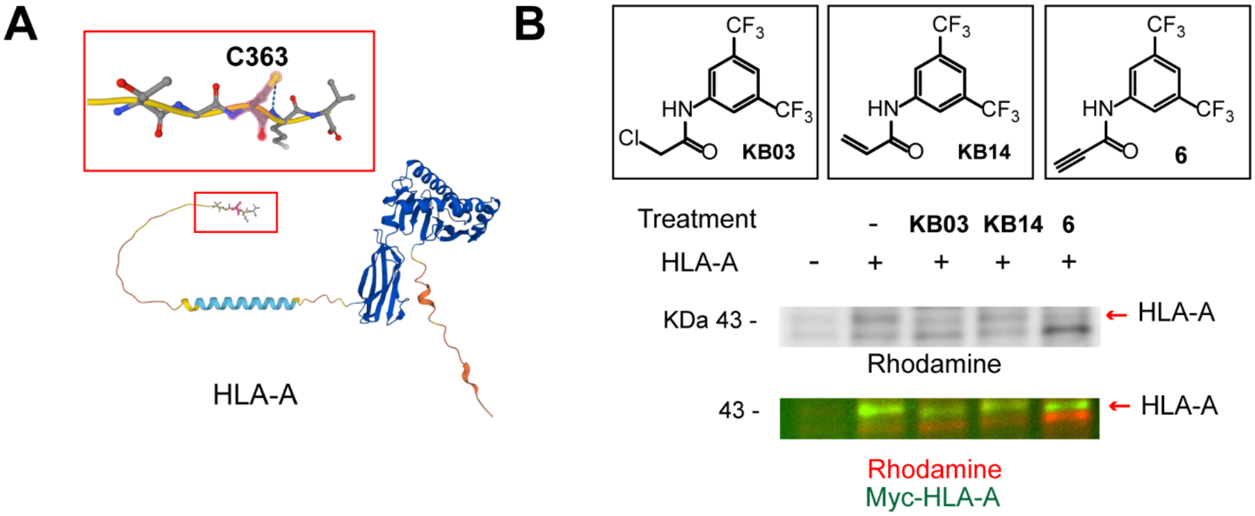


**Figure S5.** A) Predicted structure of HLA-A (AF-P04439-F1). B) IA-Rho gel and western blot of HLA-A were treated with different cysteine reactive compounds and pulled down with immunoprecipitation. Compound treatment was at 50 µM in cells for 1 h.

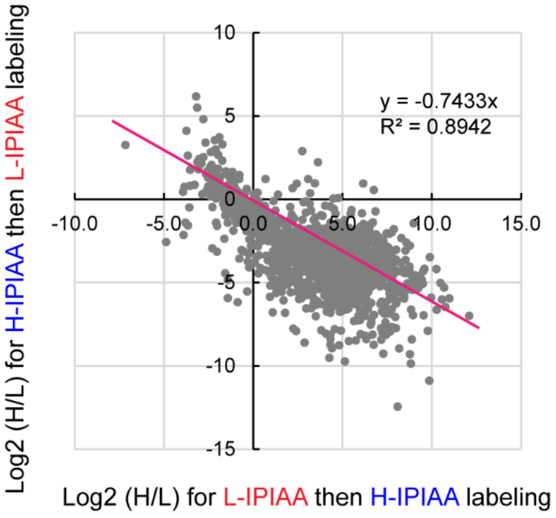


**Figure S6**. $Log_2$(H/L) reported applying L-IPIAA or H-IPIAA before reduction, respectively, when quantifying cysteine oxidation states with Cys-Surf. MS experiments were conducted in 2 replicates in Jurkats cells.

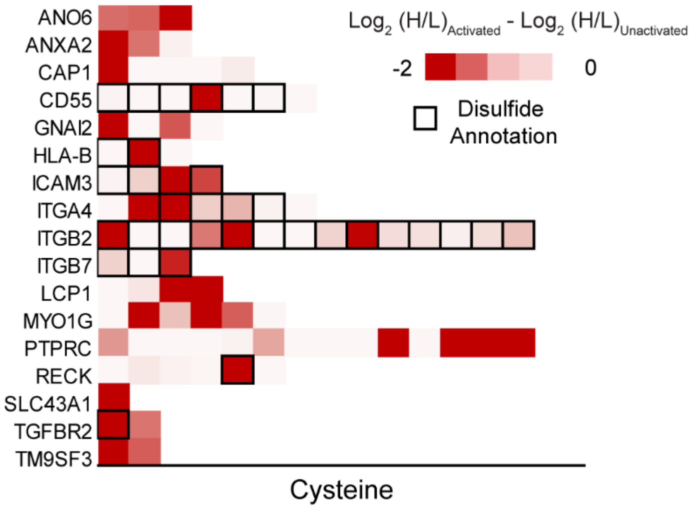


**Figure S7**. Difference of redox states for representative cysteines quantified with Cys-Surf during T cell activation. MS experiments were conducted in 3 replicates in activated and unactivated T cells.

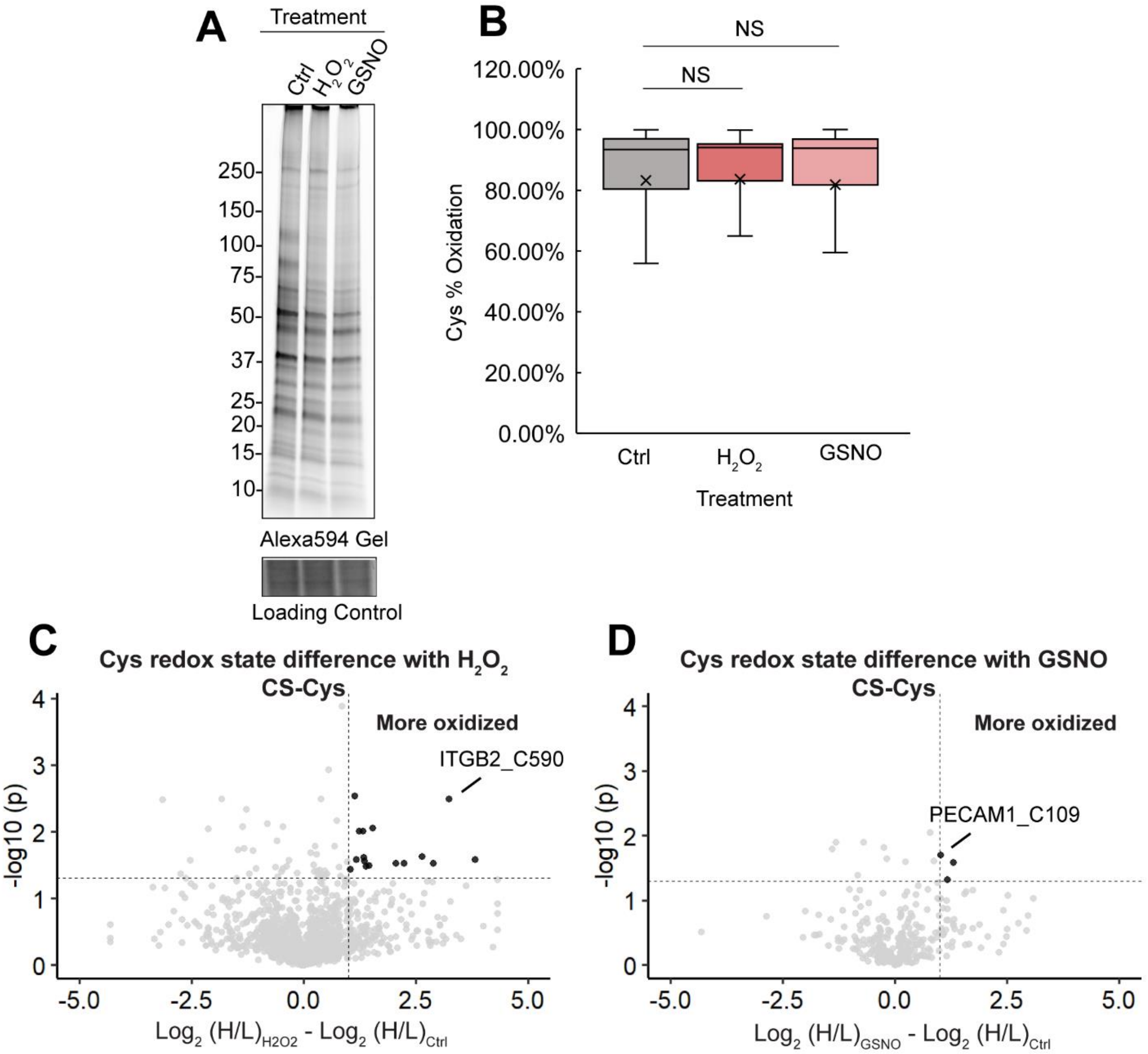


**Figure S8.** A) Gel of cell surface cysteines labeled with 5μM Maleimide-Alexa594 that were treated with DMSO as control, $H_2O_2$ or S-Nitrosoglutathione (GSNO). B) Redox states of cell surface cysteines quantified with Cys-Surf treated with $H_2O_2$, TCEP or GSNO. Statistical significance was calculated with Student's t-tests, Not Significant (NS) $p>0.05$. C) Difference of redox states for cysteines quantified with Cys-Surf with or without $H_2O_2$ treatment. D) Difference of redox states for cysteines quantified with Cys-Surf with or without GSNO treatment. For panel B, the mean is represented by "x" and the box plot is with 95% confidence interval. MS experiments were conducted in 2 replicates in Jurkats cells.

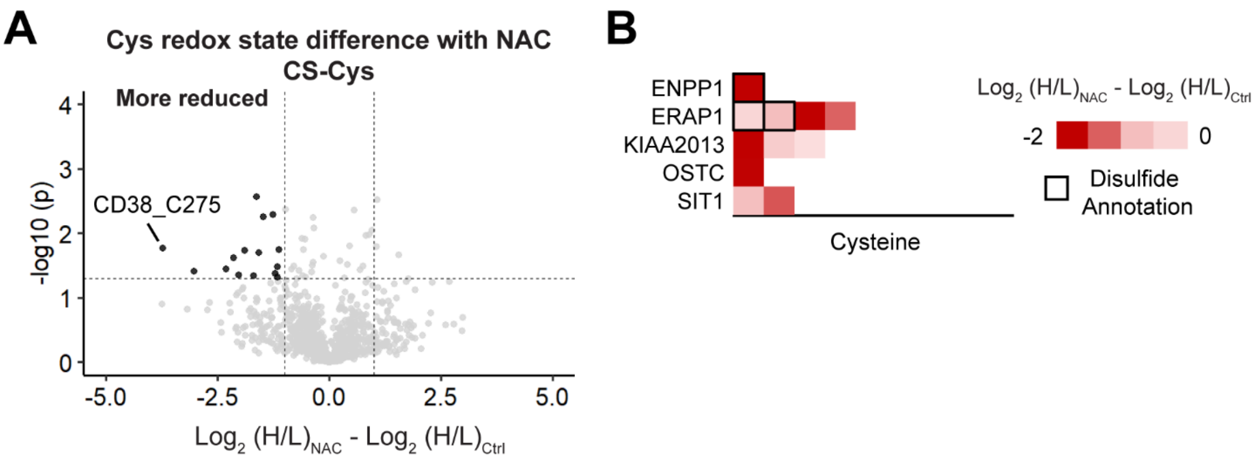


**Figure S9**. A) Difference of redox states for cysteines quantified with Cys-Surf with or without NAC treatment. B) Difference of redox states for representative cysteines quantified with Cys-Surf with or without NAC treatment. MS experiments were conducted in 3 replicates in Jurkats cells. All data can be found in **Table S5**.

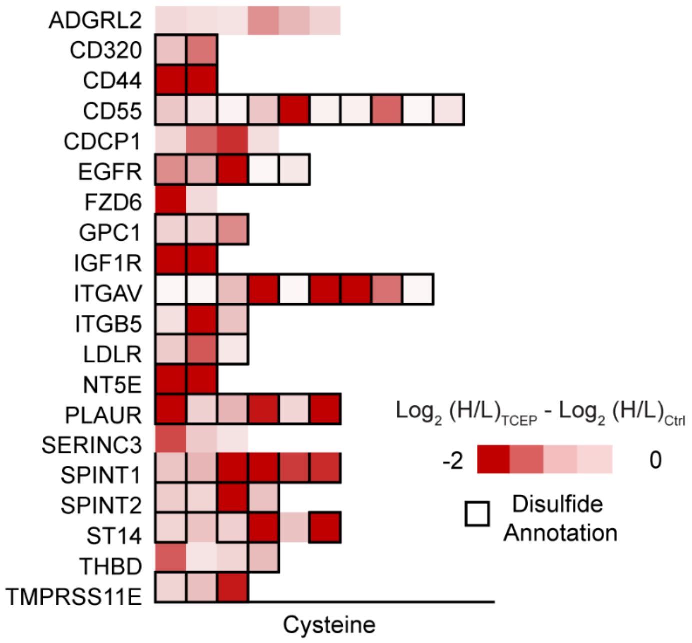


**Figure S10.** Difference of redox states for representative cysteines quantified with Cys-Surf with or without TCEP treatment. MS experiments were conducted in 3 replicates in Jurkats cells. All data can be found in **Table S5**.

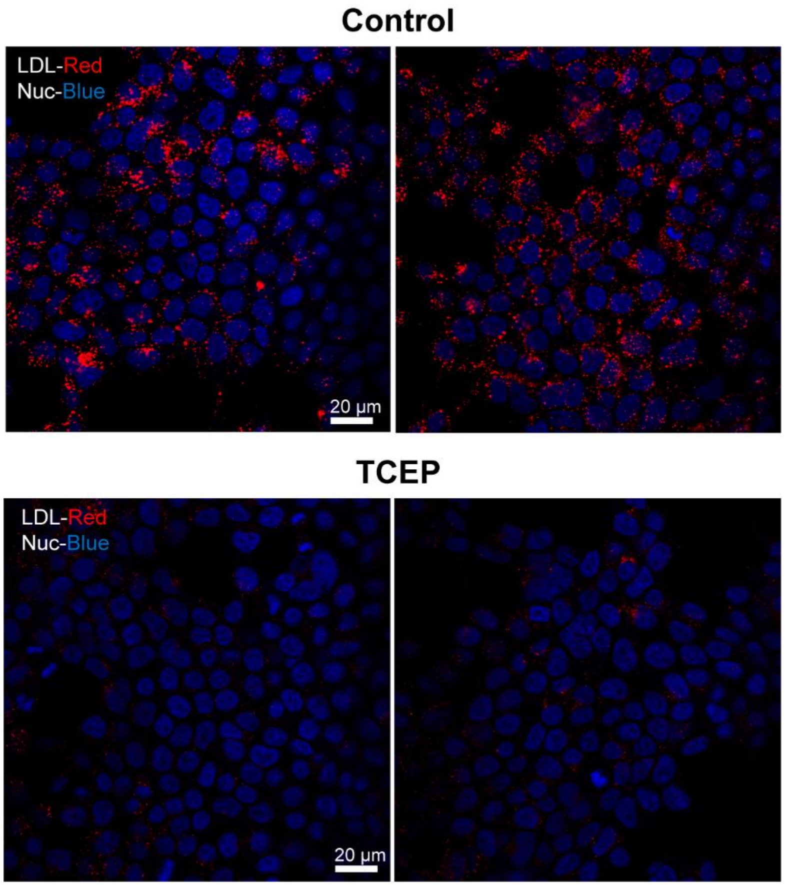


**Figure S11.** LDL uptake (signals in red) in WT HEK293T cells with or without treatment of 1 mM TCEP for 20 min. Each panel presents two different areas of cells.

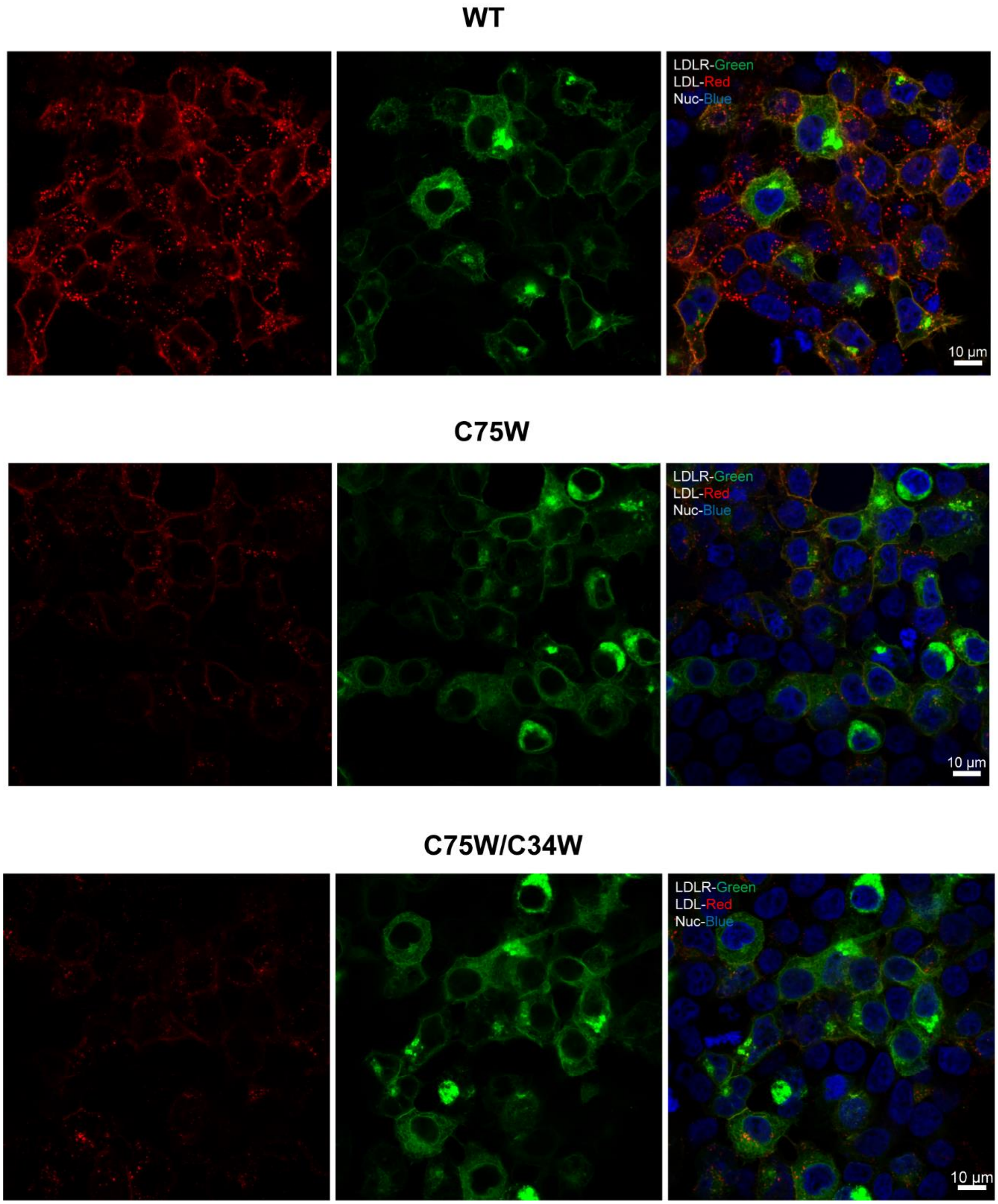


**Figure S12.** LDL uptake (signals in red) in HEK293T cells with transient expression of $LDLR^{WT}$, $LDLR^{C75W}$ and $LDLR^{C75W/C34W}$ (signals in green).

**B. Supplementary Tables**

**Table S1-5.** Datasets corresponding to each figure, provided in the attached supplementary files.

**Table S6**. Antibodies and fluorescent probes used in this study

| Name | Catalog number | Concentration |
|---|---|---|
| IA-Rhodamine | Invitrogen T6006 | 5 μM |
| Alexa Fluor™ 594 $C_5$ Maleimide | Invitrogen A10256 | 5 μM |
| IRDye® 800CW Streptavidin | LI-COR 92632230 | 1:5000 |
| Myc-Tag Mouse | Cell Signaling 2276 | 1:50 (Immunoprecipitation) |
| Myc-Tag Rabbit | Abclonal AE070 | 1:10,000 |
| IRDye® 800CW Goat anti-Rabbit | LI-COR 102673330 | 1:5000 |

**Table S7.** Conditions of Liquid-chromatography (LC)

| Parameter | Condition |
|---|---|
| Column | 100 μM ID fused silica capillary packed in-house with bulk C18 reversed phase resin (particle size, 1.9 μm; pore size, 100 Å; Dr. Maisch GmbH) |
| Mobile phase | Buffer A: water with 3% DMSO and 0.1% formic acid<br>Buffer B: 80% acetonitrile with 3% DMSO and 0.1% formic acid |
| Gradient and flow rate | 0 – 5 min, 3 – 10% B, 300 nL/min |
| | 5 – 64 min, 10 - 50% B, 220 nL/min |
| | 64 – 70 min, 40 - 95% B, 250 nL/min |
| Run time | 70 minutes |
| Injection volume | 5 uL |

**Table S8.** Files in Proteomics Identification Database (PRIDE) datasets

| Figure | File name | Experiment |
|---|---|---|
| 1 | 2021-02-01-kb-70min_FAIMS_100micron_3cv_-35_-45_-55_SY52-Try-click-40-sp3-NA<br>2021-03-18-kb-70min_FAIMS_100micron_3cv_-35_-34_-55-SY55-T5<br>2021-03-18-kb-70min_FAIMS_100micron_3cv_-35_-34_-55-SY55-T6 | Cys-Surf |
| 2 | 2021-03-05-kb-70min_FAIMS_100micron_3cv_-35_-34_-55_SY54-T1<br>2021-03-05-kb-70min_FAIMS_100micron_3cv_-35_-34_-55_SY54-T2<br>2021-03-05-kb-70min_FAIMS_100micron_3cv_-35_-34_-55_SY54-T3<br>2021-03-05-kb-70min_FAIMS_100micron_3cv_-35_-34_-55_SY54-T4 | Cys-Surf Oxidation |
| 3 | 2021-06-22-KB-SY59-3<br>2021-06-22-KB-SY59-4<br>2023-04-23-KB-SY59-3 | Cys-Surf TCEP |
| | 2021-05-30-SY-58-1<br>2021-06-03-SY-58-2<br>2021-06-22-KB-SY59-1 | Cys-Surf NAC |
| | 2021-06-22-KB-SY59-5<br>2021-06-22-KB-SY59-6<br>2021-05-30-SY-58-3 | Cys-Surf Ctrl |
| 4 | 2021-09-27-KB-SY-63-1-naive<br>2021-09-27-KB-SY-63-2-naïve<br>2023-04-23-KB-SY63-1-naive<br>2021-09-27-KB-SY-63-3-activated<br>2021-09-27-KB-SY-63-4-activated<br>2023-04-23-KB-SY63-3-activated | Cys-Surf T cell |
| 5 | 2022-03-11-KB-SY86-2<br>2022-03-11-KB-SY86-3<br>2023-04-19-KB-SY86-2 | Cys-Surf ABPP |
